# Enhancing Evolutionary Timelines: The Impact of Stratigraphic Range Information on Phylogenetic Inference

**DOI:** 10.1101/2025.04.17.649084

**Authors:** Ugnė Stolz, Alexandra Gavryushkina, Timothy G. Vaughan, Tanja Stadler, Bethany J. Allen

**Author notes:** These authors contributed equally.

## Abstract

Coherent phylogenetic analyses of molecular and fossil datasets have deepened our understanding of evolutionary biology. A core model facilitating such coherent analyses is the fossilised birth-death (FBD) model which directly incorporates fossils within phylogenetic trees. However, a limitation of the FBD model is that it cannot assign multiple fossil occurrences of the same species, limiting our ability to accurately represent age information for species which have been sampled repeatedly. To address this gap, the Stratigraphic Ranges Fossilized Birth-Death (SRFBD) model has recently been introduced. This model can account for sampled stratigraphic ranges and integrate over occurrences within the range, enabling us to include more complete fossil age information within the inference. Here, building upon this mathematical work, we develop a computational method making the model accessible to the community, for more accurate total-evidence inference of dated phylogenetic trees and evolutionary parameters. In particular, we integrate the SRFBD model into BEAST2, facilitating the use of a diverse array of genomic substitution models and the inclusion of both morphological and molecular data when inferring phylogenies. We present a thorough validation of the SRFBD implementation against simulated data. We then demonstrate the differences in posterior parameter estimates and phylogenies when applying FBD and SRFBD models to two example datasets, the Spheniscidae (penguins) and the Canidae (dogs). In both examples, the SRFBD model produces older divergence times, lower diversification and turnover rates, and considerably higher sampling proportions compared to FBD model.

## 1 Introduction

Phylogenetic trees describe the ancestral history and divergence patterns of species, and as a result, the accurate inference of phylogenies is fundamental to our understanding of evolutionary biology. The increasing availability of genomic data, combined with information from the fossil record and recent advances in Bayesian phylogenetic methods, have revolutionised the field, allowing the integration of different types of data and the estimation of evolutionary history with explicit consideration of uncertainty [1–3].

Among others, the fossilised birth-death (FBD) [4, 5] model has emerged as a coherent mechanistic framework which incorporates fossil age data directly into the analysis. The framework also allows the possibility of sampled ancestors [6], meaning that extinct species can be direct ancestors of other sampled extinct or extant species. This has significantly improved our ability to infer accurate evolutionary timelines [7].

One limitation of the FBD model is its inability to accurately include age information from a rich fossil record. Species may be known from fossils dated to multiple time intervals. The period of time over which fossils of a given species occur is known as a stratigraphic range, which provides maximum and minimum bounds on the duration of time over which the species existed [8]. When using the FBD, it is possible to include several fossils of a species, but the model is ignorant to the information that these fossils are part of the same species. In common applications of the FBD model, a species’ stratigraphic range is usually represented by a single fossil occurrence, and the range is either ignored or treated as uncertainty in the age of this single fossil [5, 9]. This violates the FBD model assumption of an equal fossil sampling rate across lineages, since lineages representing species with multiple fossil occurrences and potentially longer duration become sampled with lower intensity. Such a violation can result in underestimated sampling rates, that might in turn bias estimates of other parameters including tree topology and divergence dates. The replacement of a known stratigraphic range with a single, potentially uncertain, time point for each branch also introduces additional uncertainty in the timing of speciation and extinction, beyond that which would exist were the ranges to be included directly. This is because true stratigraphic ranges impose stronger constraints than a single fossil, since they require that speciation must occur earlier than the oldest known occurrence and extinction later than the youngest known occurrence.

To address these issues, Stadler et al. [10] introduced a stratigraphic range fossilised birth-death (SRFBD) process, incorporating three modes of speciation: asymmetric (budding), symmetric (bifurcating) and anagenetic [11]. This process models species’ durations through time and, as a result, species’ stratigraphic ranges (Figure 1). However, to date, the inference of dated phylogenetic trees and diversification parameters using the SRFBD model has not been possible due to its absence from any practical inference framework.

**Figure 1:**
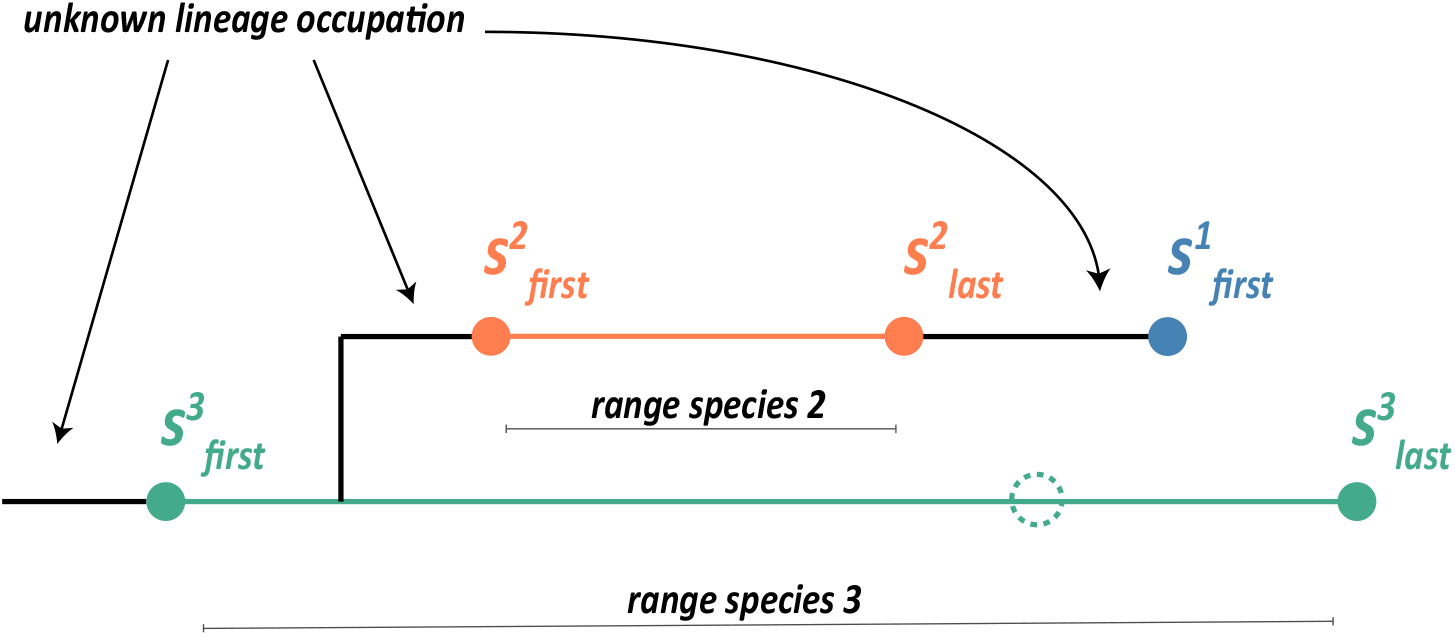
Diagram of an asymmetric speciation tree. The schematic shows three sampled species *S*^1^, *S*^2^ and *S*^3^. Species *S*^1^ has only one observation (occurrence), and therefore its known range is of length 0. Species *S*^2^ has two observations, and its range occupies the tree lineage between them. Species *S*^3^ has three observations; however, we only use the *first* and *last* occurrences to define the range, integrating over any sampling within it. The “ignored” sample is shown as a dashed circle. We have only one observed branching event, which occurs within the range of *S*^3^. Our tree is oriented and *S*^3^ is a sampled ancestor species to two sampled descendant species, *S*^1^ and *S*^2^. Since the occupation of the descendant lineage immediately after the branching event is unknown, there might have been additional intermediate descendant species, which have not been sampled. Species *S*^2^ is also a sampled ancestor of *S*^1^. Note the difference between the ancestor-descendant pair of species *S*^1^ and *S*^2^ and the pair *S*^2^ and *S*^3^. In the first case, descendant species *S*^2^ lies on the lineage that buds off the lineage of its ancestral species *S*^1^. In the second case, species *S*^2^ and *S*^1^ lie on the same “straight line” lineage, which implies that there must have been an unobserved budding speciation event on the segment of the lineage between samples 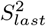 and 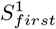.

Here, we present a framework for inferring phylogenetic trees and model parameters under the SRFBD model, using a budding mode of speciation, and report on its implementation in the BEAST2 phylogenetic inference software [1]. Our method enables Bayesian inference of dated phylogenetic trees using stratigraphic range data, in conjunction with the array of substitution and clock models already available in BEAST2. Leveraging Markov Chain Monte Carlo (MCMC) algorithm, we enable efficient exploration of the vast parameter space inherent to phylogenetic studies, aiming for accurate inference of evolutionary parameters. Our method further accounts for uncertainty in fossil ages by sampling the ages of the first and last occurrences of fossils in a stratigraphic range from intervals representing uncertainty around the ages of these specimens. This addresses the challenge that it is rarely possible to assign a precise age to fossils [12], and it has been shown that falsely forcing a precise fossil sampling date leads to bias [9].

Furthermore, our implementation of the SRFBD model produces a posterior distribution over the space of oriented trees (Figure 1), which means that at each speciation event, one of the branches is assigned to be ancestral, and the other descendant. Orientations of branching events supported by the data – such as a species descending from budding of an ancestral stratigraphic range – will receive higher posterior probabilities, and branching events without a clear orientation supported by the available data will essentially be symmetric, with equal probabilities assigned to either orientation. This allows us to test hypotheses about ancestor-descendant relationships in the tree.

In the next sections, we recapitulate the mathematical framework developed by Stadler et al. [10]. Then, we introduce our algorithm and present the results of a validation study that shows the correctness of its implementation and robustness of the underlying model. Finally, we compare FBD and SRFBD models by applying them to a combined molecular and morphological Spheniscidae (penguin) dataset, as well as Canidae (dogs) data comprised of only morphological characters.

## 2 Materials and Methods

### 2.1 Characteristics of a phylogenetic tree with ranges (SR tree)

Evolutionary relationships between species are represented by a phylogenetic tree. The incomplete nature of the fossil record, and subsequently our datasets, means that we cannot reconstruct the complete evolutionary history of species and are only able to reconstruct the observed part of the phylogenetic tree, which connects the sampled species, and is called a sampled (or reconstructed) tree [13]. This is typically a tree where each node has two or no descendants, corresponding to an internal node representing speciation or a tip node representing a sample. Lineages without descending samples are not shown in the phylogenetic tree [13]. In phylogenetic trees generated by the FBD process, there are additionally nodes with a single descendant, corresponding to sampled ancestor (SA) species, which later give rise to further sampled species [6].

When we consider stratigraphic ranges of sampled species, we need to account for different modes of speciation in our tree representation. Three main modes of speciation have been identified [14]: symmetric speciation, where an ancestral species splits into two new species and ceases to exist (so it is considered extinct); budding (or asymmetric) speciation, where an ancestral species persists while giving rise to a new species; and anagenetic speciation, where a species transforms into another species over time without branching. In symmetric and anagenetic speciation, the ancestral species does not persist and therefore its stratigraphic range must terminate at the speciation event. Under budding speciation, the ancestral species continues to exist. The divergence of a new species can occur either within the known stratigraphic range of the ancestor, or within an unsampled part of the lineage (Figure 1).

To implement these speciation modes, the tree object has to be enriched. For asymmetric speciation, we must label the two child branches as ancestor (the continuation of the old branch) and descendant (the new branch), which we visualise by consistently orientating the ancestor branch at the bottom and the descendant branch at the top (Figure 1). No such labelling is needed for symmetric and anagenetic speciation since the ancestral species does not persist in either case.

In this paper, we only consider asymmetric speciation and call an oriented tree with stratigraphic ranges a stratigraphic range (SR) tree. When there is insufficient information to determine the directionality of a speciation event, the orientation is arbitrary. Similar to the FBD case, we can identify a sampled ancestor species as a sampled species that has sampled descendants, e.g., species *S*_3_ and *S*_2_ are sampled ancestors.

### 2.2 Bayesian inference of SR trees

We seek to evaluate the posterior probability distribution:

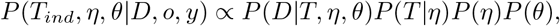

where *T* = (*T*_*ind*_, *o, y*) is a tree *T*_*ind*_ with stratigraphic ranges starting at origin time, *o*, and ending at some subsequent time *y*, generated by the SRFBD model with parameters *η* = {*λ, µ, ψ, ρ, t*_0_} (for a detailed model description see Stadler et al. [10]). We use *θ* to represent the substitution model parameters and *D* to represent the alignment data. These data could constitute molecular sequence alignments and/or a matrix describing morphological characters for the sampled species, and are attached to the nodes representing samples in the tree. For species with non-zero stratigraphic ranges, there are two nodes representing the beginning and the end of its range. We can attach the data to only one end of the range (first or last occurrence) or to both ends of the range.

Since stratigraphic range boundaries are usually not known exactly, we use *y*^*up*^ and *o*^*up*^ to represent the upper bounds on the start and ends of the stratigraphic ranges, and *y*^*low*^ and *o*^*low*^ to represent the corresponding lower bounds. To use these uncertainty intervals instead of exact ages in the inference we adjust our model as follows:

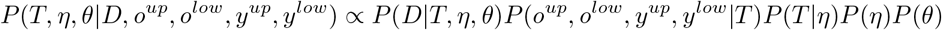

Here we do not model the procedure of assigning the age interval [*t*^*low*^, *t*^*up*^] to a fossil with true age *t* by a dating method (e.g., radiometric dating) but assume that the probability density *f* (*t*^*low*^, *t*^*up*^|*t*) does not depend on the exact position of *t* within the interval but only on whether it belongs to the interval or not. In our notation, this implies that *P* (*o*^*up*^, *o*^*low*^, *y*^*up*^, *y*^*low*^|*T*) = *P* (*o*^*up*^, *o*^*low*^, *y*^*up*^, *y*^*low*^|*o, y*) is a constant when 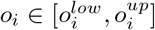 and 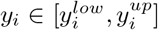 for all *i* and zero otherwise. Furthermore, as we estimate phylogeny *T* with exact ages for stratigraphic ranges, we inherently obtain estimates of the ages of the first and last occurrences of species.

To evaluate the likelihood *P* (*D*|*T, η, θ*), we use the Felsenstein pruning algorithm [15] implemented in BEAST2. We have implemented the tree prior^1^ *P* (*T* |*η*) in the *sRanges* package as described below (Section 2.2.1). Parameter prior distributions can be chosen from the variety available in BEAST2.

#### 2.2.1 Probability Density of an SR Tree

This section is a very brief summary of results from [10]. Let *η* = {*λ, µ, ψ, ρ, t*_0_}, where *λ, µ* and *ψ* are speciation, extinction, and fossil sampling rates, respectively, *ρ* is the sampling probability of extant species at the present, and *t*_0_ is the time of origin. We assume that the reconstructed tree has *n* sampled species or equivalently *n* stratigraphic ranges, as we can treat extinct species with a single fossil occurrence, or extant species without fossils, as stratigraphic ranges that start and end at the same time point. Further, we assume that *l* is the number of sampled extant species, *j* is the number of sampled ancestor stratigraphic ranges, and *k* is the total number of sampled fossils. For a stratigraphic range *i*, we denote its first and last occurrence as *o*_*i*_ and *y*_*i*_, respectively. Finally, let *I* be the set of all stratigraphic ranges *i* that lie on the same straight line as their most recent ancestral stratigraphic range (denoted as *a*(*i*)) in the tree, that is, there is no observed bifurcation event between the stratigraphic range *i* and its ancestral range *a*(*i*). All notation is summarised in table 1, which is a simplified table 1 of [10].

**Table 1:** Summary of notations.

|  |  |
| --- | --- |
| $\lambda$ | Rate of branching speciation |
| $\mu$ | Extinction rate |
| $\psi$ | Fossil sampling rate |
| $\rho$ | Extant species sampling probability |
| $t_0$ | Time of origin of a tree |
| $\eta$ | $(\lambda, \mu, \psi, \rho, t_0)$ |
| $\theta$ | Substitution model parameters |
| $d$ | Diversification rate |
| $s$ | Sampling proportion |
| $\nu$ | Turnover rate |
| $n$ | Number of sampled species (i.e., number of sampled stratigraphic ranges) |
| $l$ | Number of sampled extant species |
| $j$ | Number of sampled ancestor species |
| $k$ | Total number of sampled fossils |
| $\kappa'$ | Total number of sampled fossils representing *start/end* of stratigraphic ranges |
| $\kappa$ | Total number of fossils *within* a stratigraphic range ($k = \kappa + \kappa'$) |
| $o_i$ | Start time of stratigraphic range $i$ |
| $o_i^{up}$ | Upper bound of the age interval for $o_i$ |
| $o_i^{low}$ | Lower bound of the age interval for $o_i$ |
| $y_i$ | End time of stratigraphic range $i$ |
| $y_i^{up}$ | Upper bound of the age interval for $y_i$ |
| $y_i^{low}$ | Lower bound of the age interval for $y_i$ |
| $a(i)$ | Most recent stratigraphic range, ancestral to range $i$ |
| $I$ | Set of stratigraphic ranges, with $i \in I$ if $i$ is in the same straight line as $a(i)$ on a tree |
| $L_s$ | Length of all stratigraphic ranges on a tree |
| $D$ | Morphological and/or molecular alignment data. |
| $T$ | Reconstructed stratigraphic range phylogenetic tree |
| $T_{ind}$ | Reconstructed stratigraphic range phylogenetic tree without the ages of the stratigraphic ranges, i.e., $T = (T_{ind}, o, y)$ |

We have implemented the probability density *P* (*T* |*η*) for a sampled SRFBD tree as in Stadler et al. [10], theorem 8:

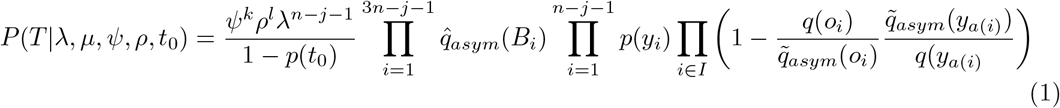

Here *p*(*t*) is the probability that an individual species alive at time *t* does not leave sampled descendants:

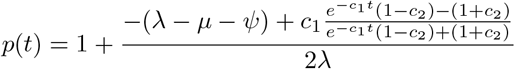

with

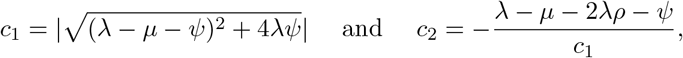

and 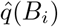 is the contribution of every tree branch *B*_*i*_ with start time *s*_*i*_ and end time *e*_*i*_:

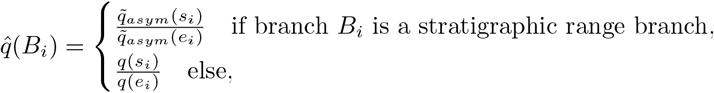

with

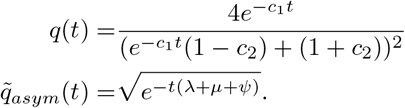

The proof is available at [10].

To avoid estimating the number of fossils within the ranges, which would be an ambiguous and time-consuming task, we also implemented the possibility to marginalise over any number of fossils observed between the first and last known samples. That is, we remove intermediate occurrences within stratigraphic ranges from the tree and instead integrate over the number of fossils within this range. This means that our estimates of *ψ*, the constant fossil sampling rate, represent sampling between and within the ranges and are not influenced by sampling frequency fluctuations between the ranges. Let *T*_*r*_ be the sampled oriented tree *T* when we ignore *κ*, occurrences sampled within the stratigraphic ranges. Then, following theorem 14 of [10]:

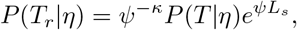

where *L*_*s*_ is the sum of all stratigraphic range lengths. Let *κ*^′^ denote the number of fossils that represent the first and last occurrences of species. Since *ψ*^*k*^ = *ψ*^(*κ*+*κ*^*′*) in the expression for *P* (*T* |*η*), the *ψ*^−*κ*^ and *ψ*^*κ*^ cancel out in the above equation and the probability density of *T*_*r*_ only depends on *κ*^′^.

### 2.3 Validation study

We performed a simulation based validation study to demonstrate that *sRanges* can accurately infer phylogenies when supplied with data, simulated under the same model. This study is inspired by the well-calibrated validation approach [16], which considers a Bayesian model well-calibrated if the highest posterior density (HPD) region at *x*% credibility level contains the true value with a frequency approximately equal to *x*%. To maintain computational feasibility, we introduce a slight difference between the simulation and inference models by conditioning our simulated trees against having excessively large or small sample sizes. Since we did not apply the same conditioning in our inference, the simulation distribution and the inference prior are not perfectly aligned. To show that this conditioning did not influence our results negatively, we also performed rank-uniformity validation (RUV, [17]), which has been shown to be sensitive to too large rejection percentages, even when not captured by the coverage analysis [18].

We simulated phylogenies under the SRFBD model, retaining only the first and last observations (fossil occurrences) of each species to constitute that species’ stratigraphic range (these might be the same observation for poorly sampled species). As our primary goal was verification of the developed inference framework, we chose parameters likely to produce highly informative input data, with fast evolutionary rates and complete morphological and molecular data alignments. The intention was to avoid overly broad posterior distributions which are computationally difficult to characterize using our MCMC algorithm. For validation, the SRFBD parameters, discussed above, were transformed to diversification rate *d* = *λ* − *µ*, turnover 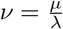 and sampling proportion 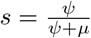.

We analysed two different data scenarios. First, we assumed maximum molecular and morphological information by attaching morphological and DNA data to both first and last stratigraphic range bounds. Second, we explored a more analytically realistic scenario by attaching morphological data to both stratigraphic range bounds, but DNA data only to extant samples. In both data scenarios, we used the same 200 simulated trees obtained as follows.

Data simulation and result analysis were performed in R [19]. Sets of parameters were drawn at random from the following distributions: *d* ∼ Uniform[0.7, 0.9], *ν* ∼ Uniform[0.2, 0.8], *s* ∼ Uniform[0.2, 0.8], *ρ* ∼ Uniform[0.7, 1]. The origin was set to *t*_0_ = 4. For each parameter draw, we simulated complete FBD trees with the *TreeSim* [20] package. Then, we used *FossilSim* [21] to simulate taxonomy for a budding speciation mode and to sub-sample the tree with a sampling proportion *s* and sampling at present probability *ρ*. Incorporating taxonomy allowed us to assign each sampled fossil to a species and derive a dataset of first and last observed species occurrences (sampled stratigraphic range bounds). We then used this dataset to simulate morphological and molecular sequences along the true trees. Morphological sequences consisted of 200 characters, each with 7 discrete states, and were simulated under the LewisMK model with a strict clock rate of 0.01 substitutions per site per unit time. Molecular sequences consisted of 1000 nucleotides and were simulated under the Jukes-Cantor (JC69) substitution model with a strict clock rate of 0.01. We rejected simulated trees with more than 1000 total samples or less than 5 extant samples, drawing new parameter sets until we reached 200 trees satisfying these conditions. In total 261 trees were rejected, of which 234 went fully extinct before the present, and the rest were rejected due to having too many or too few samples.

Inference was conducted using the developed *sRanges* package, setting prior distributions to the same distributions that we used for simulating the data. For the origin, we used a Uniform[1, 1000] prior distribution. We used the true sample times and simulated sequences as input data. We specified the true evolutionary models and parameters for both DNA and morphological data. The output was analysed and plotted using R [19] packages *stringr* [22], *coda* [23] and *ggplot2* [24], *ape* [25], *beastio* [26], *R*.*utils* [27]. The RUV test [17] was performed using *SBC* [28] package, following the procedure in. See Supplementary Material for more information on code availability.

### 2.4 Penguin and Canid data analysis

To investigate SRFBD performance using real datasets, particularly in comparison with the FBD which cannot account for stratigraphic ranges, we inferred phylogenies for two families of vertebrates, namely penguins (Spheniscidae) and dogs (Canidae). In all data analyses, we used the same parameter configuration as in the simulation studies (Section 2.3). Time is measured in million years before the present (Ma).

#### Penguin data

The evolution of penguins (family Spheniscidae) is noteworthy for several reasons. Firstly, the group has an unusual ecology, being flightless birds with a semi-aquatic life habit [29, 30]. Secondly, fluctuations in the diversity and paleogeography of the clade over time have been linked to changes in climate [29–31], and the formation of islands [32], in the Southern Hemisphere over the Cenozoic. Dramatic changes in body size have also occurred across the clade [33–35]. Thirdly, the clade has a relatively rich fossil record, but of morphologies fairly distinct from extant penguins. This has led researchers to speculate previously about the relationship between the crown (living penguins and all extinct species nested within this clade) and the stem (extinct penguin clades which predate living species) [5, 31, 33]. Fourthly, there is also a high availability of molecular sequence data for penguins, enabling the integration of both morphological and molecular evolution into phylogenetic analyses [5, 30, 33]. However, in spite of this data availability, previous studies have inferred somewhat conflicting timelines for the evolution of penguin diversity.

We analysed datasets containing 55 penguin (*Spheniscidae*) species, of which 36 are extinct and 19 are extant, as in Gavryushkina et al. [5]. (While datasets that include more recently discovered species exist [34, 36], the major focus of this paper is model comparison, and hence we chose a dataset that was analysed with the FBD model before.) Morphological data, comprised of 264 characters, was available for all 55 species [37]. Characters had between 2 and 7 states, with the majority (192) having 2 states (this also includes 48 characters with binary ”present/absent” coding). Between 0.78% and 97.44% character states were unknown for each species (median at 71.21%, Supplementary Table A.5). DNA sequences (for the genes RAG-1, 12S, 16S, COI, and Cytochrome B) for the 19 extant species were available, with a total concatenated sequence length of 8145 positions [37]. Between 21.5% and 86.2% of the concatenated alignment positions had unknown values for each species (Supplementary Table A.5). Fossil occurrence data were compiled from the Paleobiology Database [38], with additional cleaning conducted to remove uncertain occurrences, update occurrence ages with respect to recent literature, and align identifications with the species names included in our datasets (see the Supplementary file Penguin dates.xlsx). These age data were used to define the stratigraphic ranges for each species. 22 species (of which 5 are extant species with an existing fossil record) are represented by more than one temporally distinct occurrence in the fossil record (defined as at least half a million years apart), and hence can be attributed a stratigraphic range. See Data and Software Availability for information on example BEAST2 XMLs for these analyses.

We ran four analyses for the penguin dataset. Each analysis was run in three independent repetitions with different random starting states. The convergence of each chain was checked using Tracer [39], confirming that the effective sample size (effectively independent draws from the posterior in the chain; ESS) values for parameters and posterior were at least 200 for each chain and that all chains converged to the same empirical posterior distribution.

##### FBD, old dates (FBD_old_)

An FBD model run as in Gavryushkina et al. [5], i.e., a single fossil age was sampled for each species from a uniform prior covering the full age range (including uncertainty) of known fossils, using the age data from this previous study. This analysis exactly replicates that conducted by Gavryushkina et al. [5]. Each chain length was 10^8^, sampling every 10000, producing 9000 samples in total. The chains were combined, after discarding 10% burn-in, resulting in 27000 samples.

##### FBD updated dates (FBD)

An FBD model run as above, but with our updated fossil age data. Each chain length was 10^8^, sampling every 10000, producing 9000 samples in total. The chains were combined, after discarding 10% burn-in, resulting in 27000 samples.

##### SRFBD, updated dates, morph. data at the first occurrence (SRFBD_first_)

An SRFBD model, incorporating stratigraphic ranges spanning the first and (in the case of multiple) last samples of each species, according to the updated fossil age data. Fossil dates were sampled from a uniform prior distribution spanning the possible ages (including age uncertainty) of each fossil occurrence. Morphological data were attached to the first occurrence of each species; this assumes that the physical characteristics defining the species can all be observed in this oldest fossil, but does not make any assumptions about the subsequent morphological evolution of the species. Each chain length was 3 × 10^8^, sampling every 5 × 10^4^, 6000 samples in total. The chains were combined, after discarding 10% burn-in, resulting in 16200 samples.

##### SRFBD, updated dates, morph. data at both ends (SRFBD_both_)

An SRFBD model, incorporating stratigraphic ranges spanning the first and (in the case of multiple) last samples of each species. Fossil dates were sampled from a uniform prior distribution spanning the updated possible age interval of each fossil occurrence. Morphological data were attached to both the first and last occurrences of each species; this assumes that the physical characteristics defining the species had arisen prior to the first occurrence, and did not change throughout the time over which the species was sampled. Each chain length was 2×10^8^. To obtain the same number of samples, we sub-sampled the original analyses, sampling every 33333, 6000 samples in total. The chains were combined, after discarding 10% burn-in, resulting in 16200 samples.

The prior parameter distributions were the same for all four analyses, and match those of the previous FBD study [5]: *t*_0_ ∼ Uniform(0, 160), *d* ∼ LogNormal(−3.5, 1.5), *ν* ∼ Uniform(0, 1), *s* ∼, Uniform(0, 1), *ρ* = 1. These priors allow for large uncertainty in the evolutionary processes and fossil sampling schemes, but assume that every extant taxon was observed. As in the original analysis, we used uncorrelated relaxed log-normal clocks [40] separately for morphological and molecular data. The Lewis MKv [41] evolutionary model was used for morphological character evolution, split into six partitions by the number of states. The general time reversible (GTR+Γ) model [42, 43] was used for molecular character evolution.

#### Canid data

The evolution of dogs (family Canidae) has also been a popular but controversial topic of study. A wide range of data types is available for establishing their evolutionary history, including molecular sequences and coding of their morphological, developmental, ecological and behavioural characteristics [44–46]. However, canid morphological datasets are often heavily dependent on cranial characters (because these are most commonly fossilised), meaning that homoplasy due to similar diets can be problematic [46]. The clade has a wide geographic spread, with the colonisation of new continents over time linked to diversification [47]. The canid fossil record is particularly rich in North America, and many studies of their evolutionary history have focused on this continent [48–50]. These studies reveal abundant convergent evolution based on diet [48], and subsequent high levels of interspecific competition [49, 50].

Here, we analysed a fossil rich dataset of Canidae with morphological data (123 characters, having between two and five states, with 30.9% missing data) for five extant and 116 extinct species of North American canids, as in Slater [48]. Similarly to the penguin dataset, fossil age data was drawn from the Paleobiology Database [38], then cleaned and updated to verify included taxonomic and temporal information. 104 species have stratigraphic ranges with first and last occurrences, while 17 species are represented by a single fossil occurrence. We analysed this dataset with four model configurations:

##### FBD, old dates fixed (FBD_old_)

In this analysis, we used the same fossil age data as in [48], and the ages were fixed. We used the FBD model and estimated *ρ*. The five extant species (*Canis latrans, Canis lupus, Cuon alpinus, Lycaon pictus, Urocyon cinereoargenteus*) had ages fixed to zero and were assumed to be *ρ*-sampled. Chain length 200M, sample every 10000, 18000 samples in total after removing burn in.

##### FBD *ρ* = 0, new dates with ranges (FBD)

An FBD model with updated range information for all species. The ages of the fossil samples were uniformly sampled between the oldest estimate for the first occurrence and youngest estimate of the last occurrence. Here, *ρ* was set to zero to reflect a scenario in which extant species are included in the analysis based on having fossil samples, as opposed to being randomly sampled at present. The tree was offset (an additional parameter was added to describe the age of the youngest fossil sample, and hence the relationship between the age of the tree and the present day at time zero). The lower bound for the five extant species was set to zero. Chain length 100M, sample every 10000, 9000 samples in total after removing burn in.

##### SRFBD, new dates, morph. data at the first occurrence (SRFBD_first_)

An SRFBD model with updated stratigraphic ranges for all 121 species, and age ranges for extant species finishing at time zero. The ages of the fossil occurrences were sampled within the estimated age boundaries. *ρ* was estimated. The morphological data was attached to the first occurrence of each species. Chain length 5000M, sample every 100000, 45000 samples in total after removing burn in.

##### SRFBD, new dates, morph. data at both ends (SRFBD_both_)

The same as in the previous analysis, but with the morphological data attached to both the first and the last occurrences of each species. Chain length 1000M, sample every 10000, 90000 samples in total after removing burn in.

We set uniform prior distributions (improper in the case of unbounded parameters) for all FBD model parameters: *t*_0_, *d, ν, s*, and *ρ*. We chose an uncorrelated relaxed log-normal clock model [40] and the Lewis MK [41] evolutionary model for morphological data, split into four partitions by the number of states (two, three, four or five).

FBD and SRFBD models assume that both *ρ* and *ψ* sampling occur with equal probability among all coexisting lineages. However here, we analysed only North American canids and only species that were originally analysed in [48]. Although the nature of *ψ*-sampling incorporates stochasticity in the fossilisation, fossil discovery and species selection processes (similarly for *ρ*-sampling), including species based on their geographical location could potentially affect the results of our analyses. We hypothesize that the potential biased sampling would only have a minor effect on the results but more research should be done in this direction.

### 2.5 FBD and SRFBD clade set comparison

Most tree inference methods produce strictly bifurcating, non-oriented trees, and hence most of the tools designed for summarising and comparing tree topologies are suited to such trees. However, the modelling of speciation as a budding process and the inclusion of inferred stratigraphic ranges means that SRFBD trees are not straightforward to summarise or interpret. To address this, we adjusted the tree comparison tool originally developed for the BEAST2 package *Babel* [51], which is designed for inferring language evolution. Our version of this tool takes posterior samples of the species trees produced by the FBD and SRFBD models as input, and compares the posterior support and heights for every clade (group of species) estimated by both methods. In sampled ancestor trees generated by the FBD model, and stratigraphic range trees generated by the SRFBD model, the most recent common ancestor (MRCA) of a group of species might be sampled, leading to ambiguity regarding the time point that represents the origin of that clade. For sampled ancestor trees we follow [5]: when a sampled ancestor is the MRCA of the remaining members of the clade, we define the time of origin of the clade as the age of the sampled ancestor. In a stratigraphic range tree, divergences can occur within stratigraphic ranges, and so the MRCA of a clade can originate within the stratigraphic range of its parent species (Figure 3). In this case, we assume that the origin time of the clade coincides with the time of the divergence and the portion of the range older than the speciation event does not belong to it. This is a more relaxed, but also more informative, definition than one requiring the whole species range to be within a clade, as otherwise nested clades will have the same age in the case when several species diverge within a stratigraphic range of the same species, artificially pushing the age of the nested clade to be older as compared to the same situation in an FBD analysis.

## 3 Results

### 3.1 Inference of speciation parameters

The results of the validation study when molecular data was included for extant samples only are shown in Figure 2. All parameters are well recovered: the highest posterior density (HPD) interval width is approximately equal to the fraction of times the true value falls within it. Supplementary Figure A.10 shows that this is also true for the analysis when molecular data was included for all bounds of stratigraphic ranges. Supplementary Figures A.11, A.12 show that validation also passes the rank-uniformity test. These results mean that our method can successfully infer phylogenetic model parameters in macroevolutionary settings.

**Figure 2:**
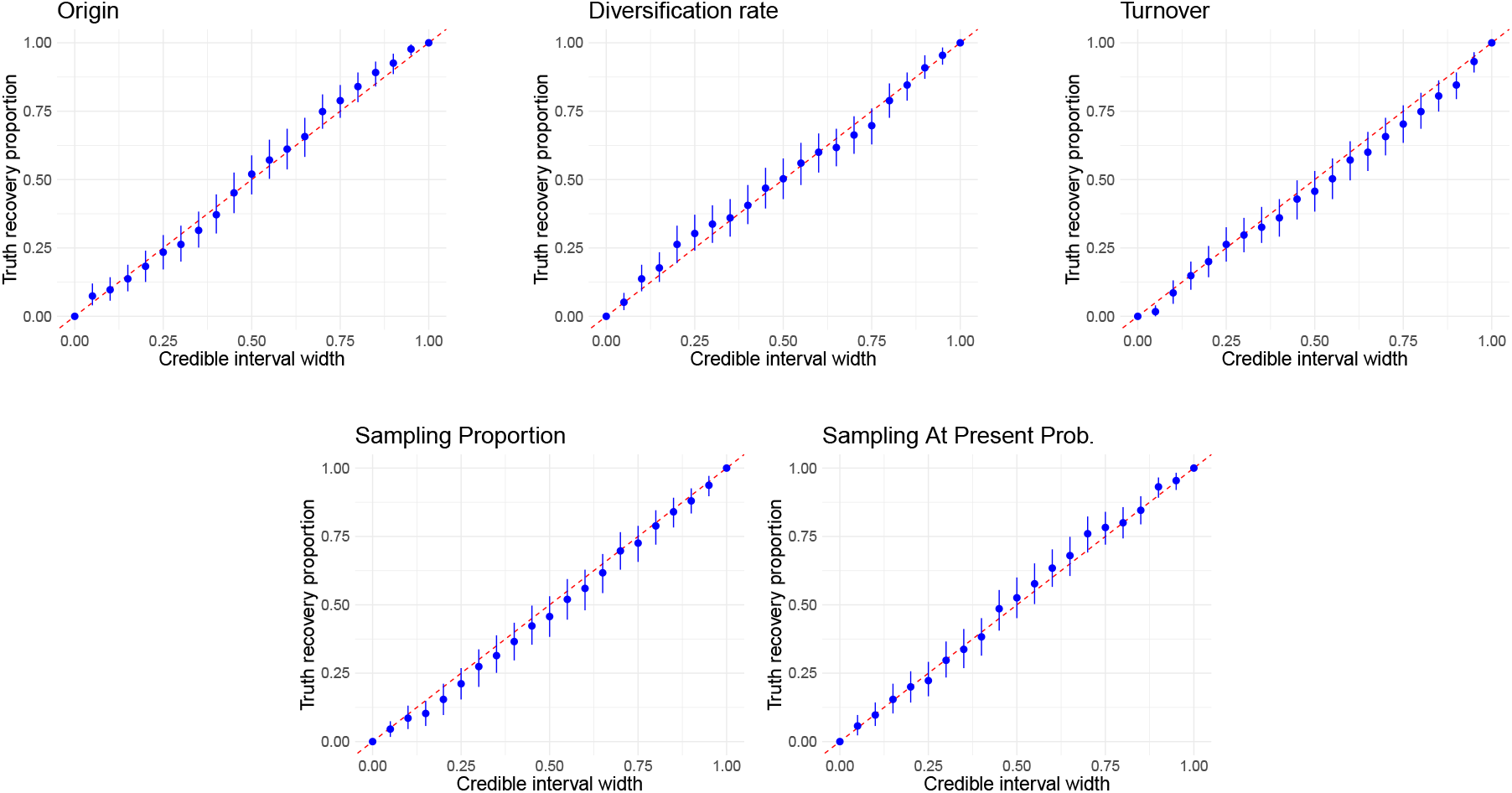
Results of the validation simulation study where molecular data is attached to extant samples only. Blue dots show the fraction of analyses where the true value is within the HPD credible interval of a certain width. The blue vertical lines show bootstrap confidence intervals. The diagonal red dashed line *x* = *y* is the expected ideal result.

**Figure 3:**
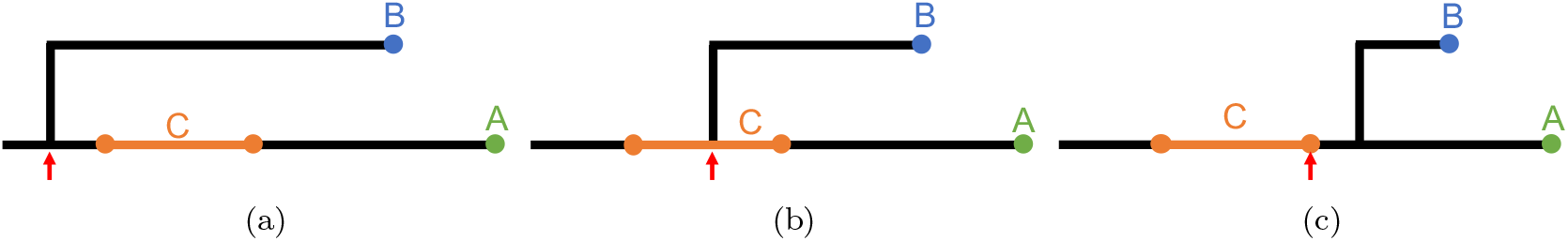
Illustration of a clade in *sRanges* package. In all three cases, we have two species A and B, represented by a single sample range (length 0), and a third species C represented by a range with two distinct occurrences (length *>* 0). We illustrate the three possible root locations of a clade that consists of all three species. That is, clade root is older than the whole range C (a); clade is rooted within the range C (b); clade is rooted at the last occurrence of species C if the clade (A, B) is rooted more recently. Note that the orientation of the clade root child branches could be reversed in cases (a) and (c), without a change in clade definition.

### 3.2 Stratigraphic ranges alter speciation inference

We compared the outputs of the FBD and SRFBD models when applied to the penguin and canid datasets in a number of ways, to cover the broad spectrum of research questions which might be addressed by inferring a time-scaled macroevolutionary phylogeny [7]. These include comparison of the tree-generating diversification and evolutionary rate parameter distributions, as well as divergence dates and topological differences. Unless otherwise specified, in this section by “FBD” we always mean the model with the updated fossil ages.

#### Penguin data

For the penguin dataset, the SRFBD obtains median posterior estimates similar to those of the FBD model for tree statistics, diversification and turnover rates (Table A.6). Parameter uncertainty was generally reduced in the SRFBD posterior distributions, with narrower 95% HPD intervals than those produced using the FBD model (Figure 4(a)-(c), Supplementary Table A.6). The sampling proportion is much higher in SRFBD analyses: median value for the SRFBD model with morphological data at first occurrence only was 0.42 (95% HPD [0.15, 0.57]), while it was 0.25 (95% HPD [0.08, 0.48]) for the FBD. Once we account for the ratio of total known occurrences for extinct taxa (69) and fossils used in the FBD analysis (36), we get the adjusted FBD sampling proportion of 0.48, 95% HPD[0.15, 0.57], which is very close to the SRFBD estimate.

**Figure 4:**
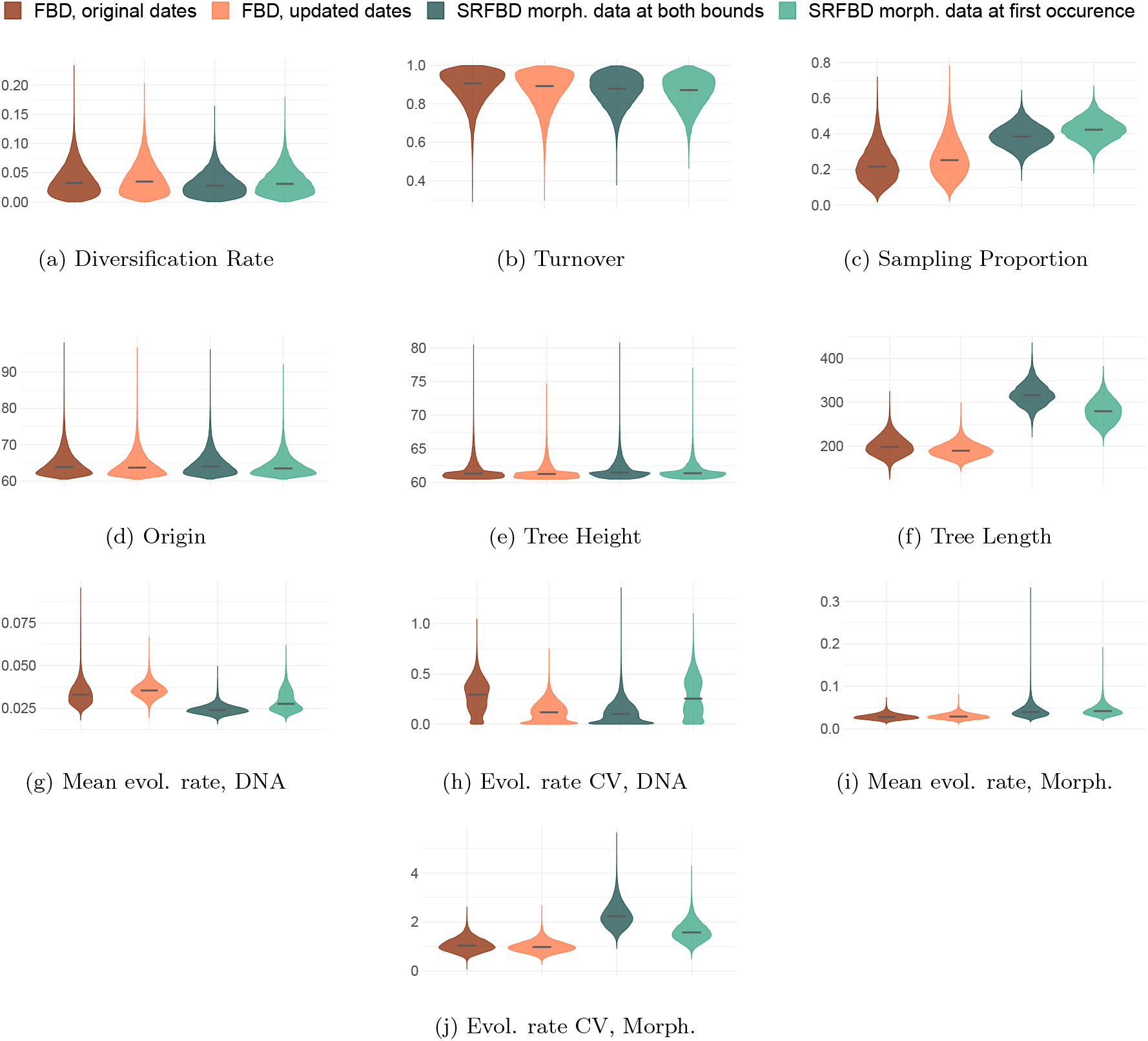
Penguin data. Parameter estimates and tree statistics obtained using the FBD, the FBD with updated dates and two SRFBD models, when attaching morphological data to either first occurrence (start of the species range) or both first and last occurrence (both bounds of the species range). The SRFBD model always used updated dates. The black line shows the median. CV = coefficient of variation.

Although the medians and 95% HPD intervals of mean per-branch evolutionary rates for molecular data are comparable between models (Supplementary Table A.6), the SRFBD with morphological data attached at first occurrence only, recovers a slightly bimodal distribution (Figure 4(g)). The minor mode agrees with the median estimate produced by the FBD model, while the major mode agrees with estimates from the SRFBD model with morphological data at both bounds. The SRFBD model recovers higher uncertainty in mean evolutionary rates for morphological data (Figure 4(i), Supplementary Table A.6) and recovers their median value to be higher than that of the FBD.

The number of species inferred to be ancestral to other sampled species estimated by the two methods differs, with the SRFBD estimating a median of 13 sampled ancestors, and the FBD a median of 19 sampled ancestor species. However, the 95% HPD intervals partly overlap (Table 2, Figure 5(a)). A clear example is *Spheniscus humboldti*, which has a SA probability of 0 in FBD, and 1.0 in SRFBD analyses (Figure 5(a)). Overall, the two methods return highly inconsistent results, with 17 species having SA probability higher or equal to 0.5 (more likely than not) in FBD, but lower than 0.5 in SRFBD analyses. The reverse is true for 7 species.

**Table 2:**
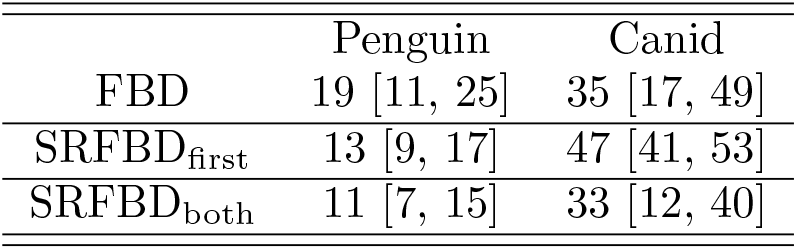
Median number of sampled ancestor species, [95% HPD]

|  | Penguin | Canid |
| --- | --- | --- |
| FBD | 19 [11, 25] | 35 [17, 49] |
| SRFBD<sub>first</sub> | 13 [9, 17] | 47 [41, 53] |
| SRFBD<sub>both</sub> | 11 [7, 15] | 33 [12, 40] |

**Figure 5:**
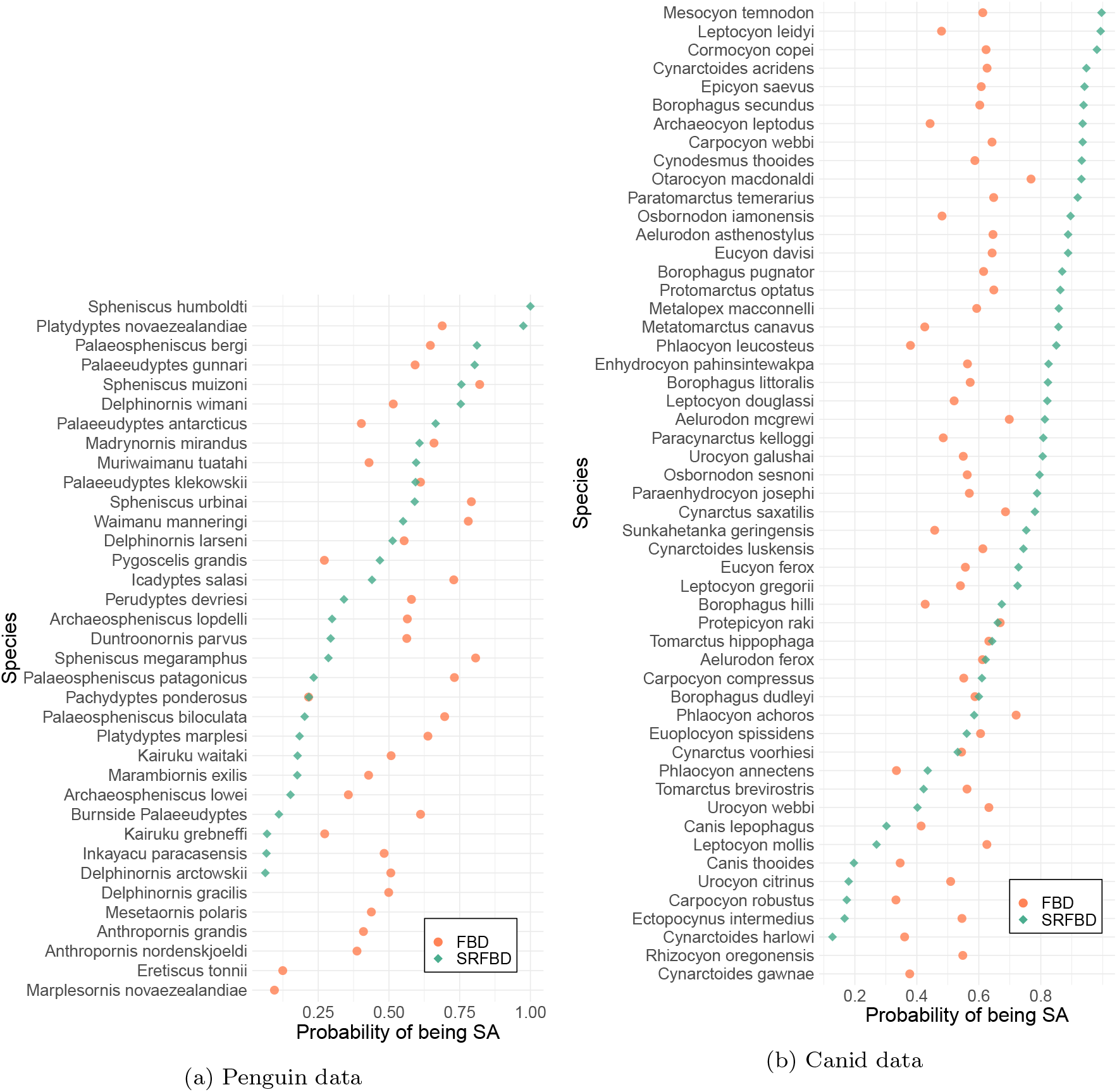
Posterior probability of species being a sampled ancestor. Circles for results of the FBD analysis, and diamonds for results of the SRFBD_first_ analysis. (a) Only species with probabilities of being a SA higher than 0.05 with either model are included. (b) Only species with probabilities of being a SA higher than 0.3 with either model included (see Supplementary Figure A.14 for the full figure).

We compared the posterior ages and support of each unique clade estimated under the FBD and SRFBD models (Figure 7 (a) and (b)) (see *FBD and SRFBD clade set comparison* for methodological context). Since the results are consistent between SRFBD_first_ and SRFBD_both_, we next compare only the SRFBD_first_ and FBD in detail. Out of 4437 unique clades obtained by either model, we further analyse 80 clades with posterior support from either method greater than 0.2. Of these, on average, the SRFBD model estimates clade ages to be 1.6 Ma older than the FBD model. For almost all (73 out of 80) clades, SRFBD estimated older ages, with no or incomplete 95% HPD interval overlap with FBD (Supplementary Figure A.15). However, the age of the tree root (tree height) is estimated to be nearly identical by both SRFBD and FBD (Figure 4 (e), Supplementary Table A.6).

The mean difference in clade support probability between the models was 0.14, and we identified 30 clades with this difference greater than the mean. The maximum difference in posterior support between the two models was 0.58, for a clade including the genera *Eudyptes–Megadyptes–Eudyptula-Spheniscus* (SRFBD 0.01, FBD 0.59). In particular, two species had very different placements in the tree between the two models (Figure 6). *Marplesornis novaezealandiae* is placed within the *Spheniscus* clade with a probability of 0.55 by the SRFBD model, versus a probability of 0.28 by the FBD model. *Madrynornis mirandus* is placed as ancestor or sibling to the *Eudyptes-Megadyptes* clade with a probability of 0.61 by the SRFBD model, versus 0.04 by the FBD model. The posterior value for *M. mirandus* to be a sampled ancestor of this clade was 0.14 in the original analyses [5].

**Figure 6:**
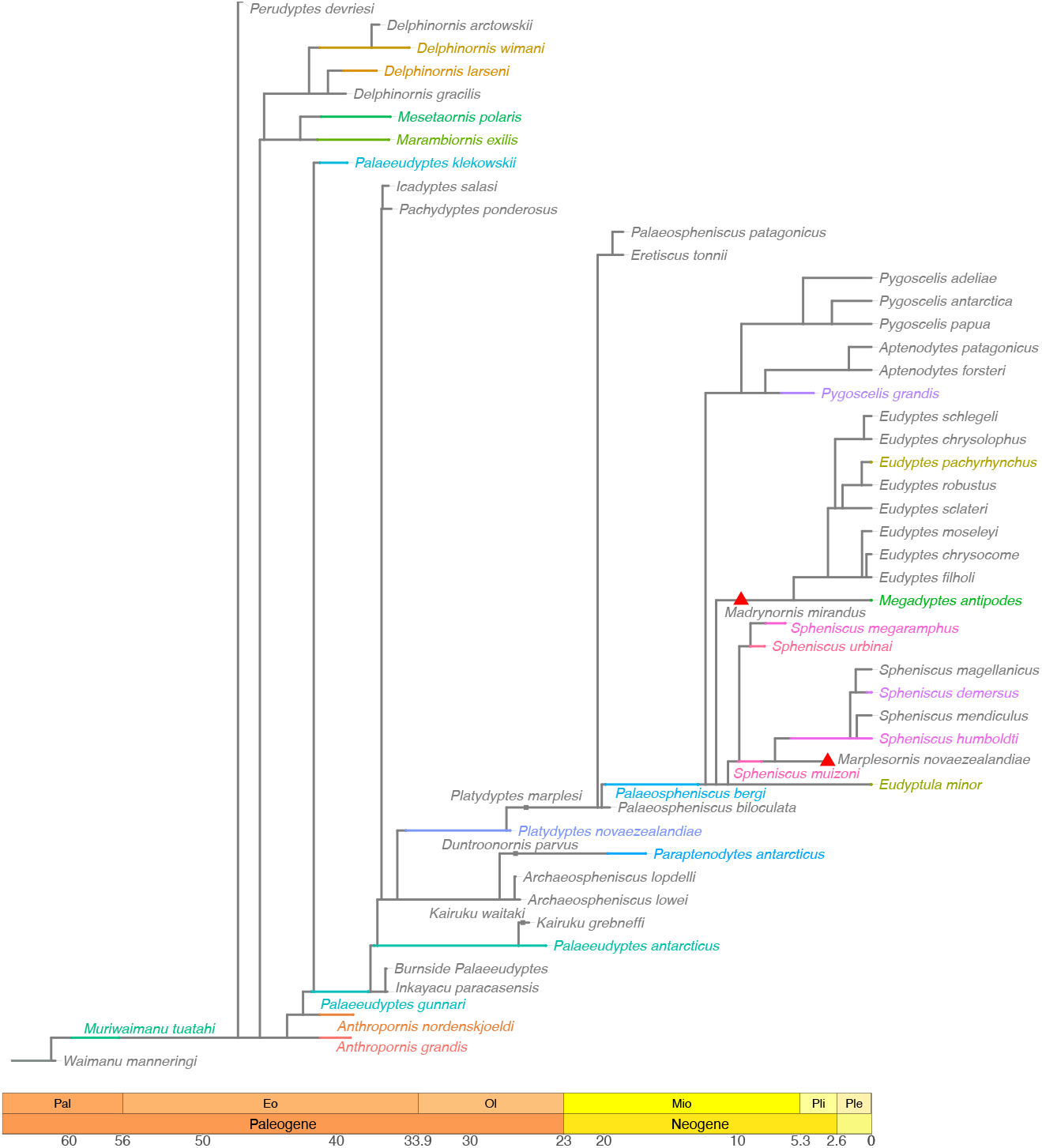
Tree with the maximum posterior probability value for penguin data. Ranges with distinct start and end points are colored. Sampled ancestor species are marked by square points. Red triangles mark *M. mirandus* and *M. novaezealandiae* species. Time in millions of years before the present (Ma).

**Figure 7:**
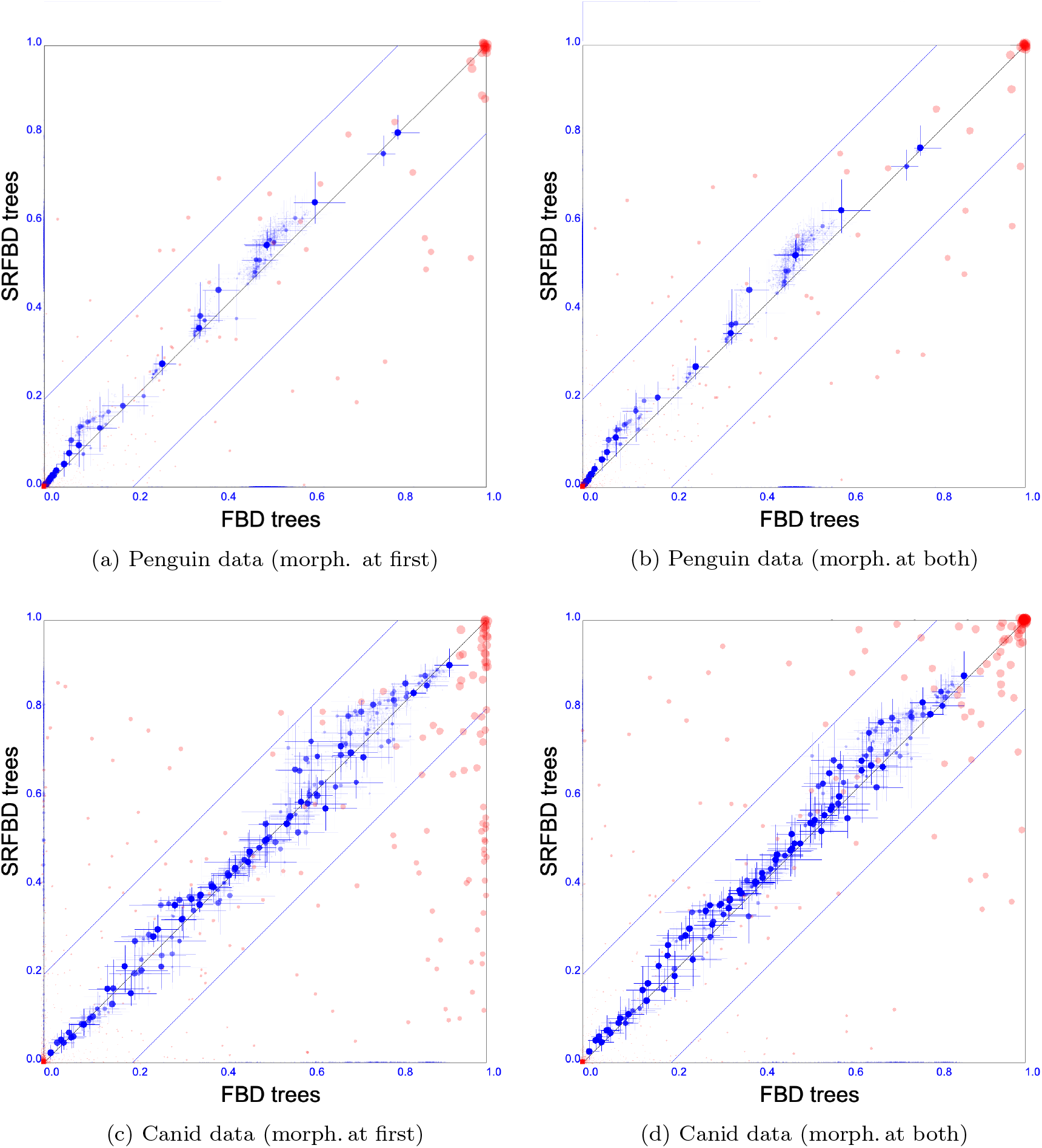
Clade set comparison between FBD and SRFBD trees. Red dots show posterior clade support, and their size shows the combined clade support. Blue dots show mean clade heights (scaled by highest corresponding tree root), while the lines show 95% HPD intervals. The clade definition for SRFBD trees is given in the Methods section.

Furthermore, both species are estimated to be part of the crown clade with higher probability by the SRFBD model (Table 4). This may be an effect of incorporating the ranges of *Spheniscus humbolti, Spheniscus urbinai, Spheniscus megaramphus, Spheniscus muizoni, Pygoscelis grandis* species within the crown: including these ranges pushes the root of this clade, on average, back by 3.07 Ma (Table 3). This topological uncertainty is particularly interesting for *M. Novaezealandiae* – there has been no consensus previously as to whether this clade is part of the crown. However, it shares several morphological characteristics with known crown species [52]. Additionally, the SRFBD cannot rule out that the crown clade is rooted within the *Palaeospheniscus bergi* stratigraphic range (posterior probability of 0.3).

**Table 3:**
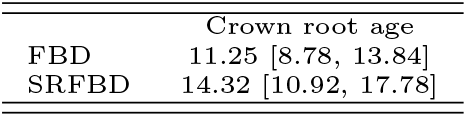
The root age of the penguin crown clade in Ma. 95% HPD interval within the square brackets.

**Table 4:** Probabilities for extinct species to be part of the penguin crown clade. Species names are **bold** if difference between the models is greater than 0.2.

| Species | FBD | SRFBD |
| --- | --- | --- |
| *Spheniscus megaramphus* | 1 | 0.99 |
| *Spheniscus urbinai* | 1 | 0.99 |
| *Spheniscus muizoni* | 1 | 0.99 |
| *Pygoscelis grandis* | 0.98 | 1 |
| *Madrynornis mirandus* | 0.79 | 0.99 |
| *Marplesornis novaezealandiae* | 0.32 | 0.71 |
| *Palaeospheniscus bergi* * | 0 | 0.3 |

\*Whenever *P. bergi* is assigned to the crown clade by the SRFBD, its first occurrence is older than the clade root and the root is within *P. bergi* range.

Finally, while sampling ages for both bounds of a species range may increase uncertainty in clade age estimates and is computationally costly, the posterior age distributions obtained by the SRFBD model are generally more informative than known uniform dating intervals, with more pronounced differences for first occurrences (Supplementary Figure A.13a).

#### Canid data

For the canids, we obtain generally similar trends as observed for the penguins. Specifically, while posterior parameter distributions largely overlap (Figure 8 (a) and (b) and Supplementary Table A.7), the median estimated diversification rate decreases from 0.073 under FBD model to 0.045 under SRFBD_first_, and median estimated turnover rate decreases from 0.94 under FBD model to 0.85 under SRFBD_first_. The sampling proportion is again much higher for the SRFBD_first_ analysis, with the median estimate of 0.78, than for the FBD analysis with the median of 0.12 (Figure 8(c)). Adjusting the FBD estimate by the ratio of known (369, based on entries with distinct ages in the Paleobiology Database) versus included (116) fossils of N. America canids, we get the sampling proportion median of 0.38, with 95% HPD interval of [0.03, 0.83]. While the uncertainty interval is very wide, it now includes the median SRFBD estimate.

**Figure 8:**
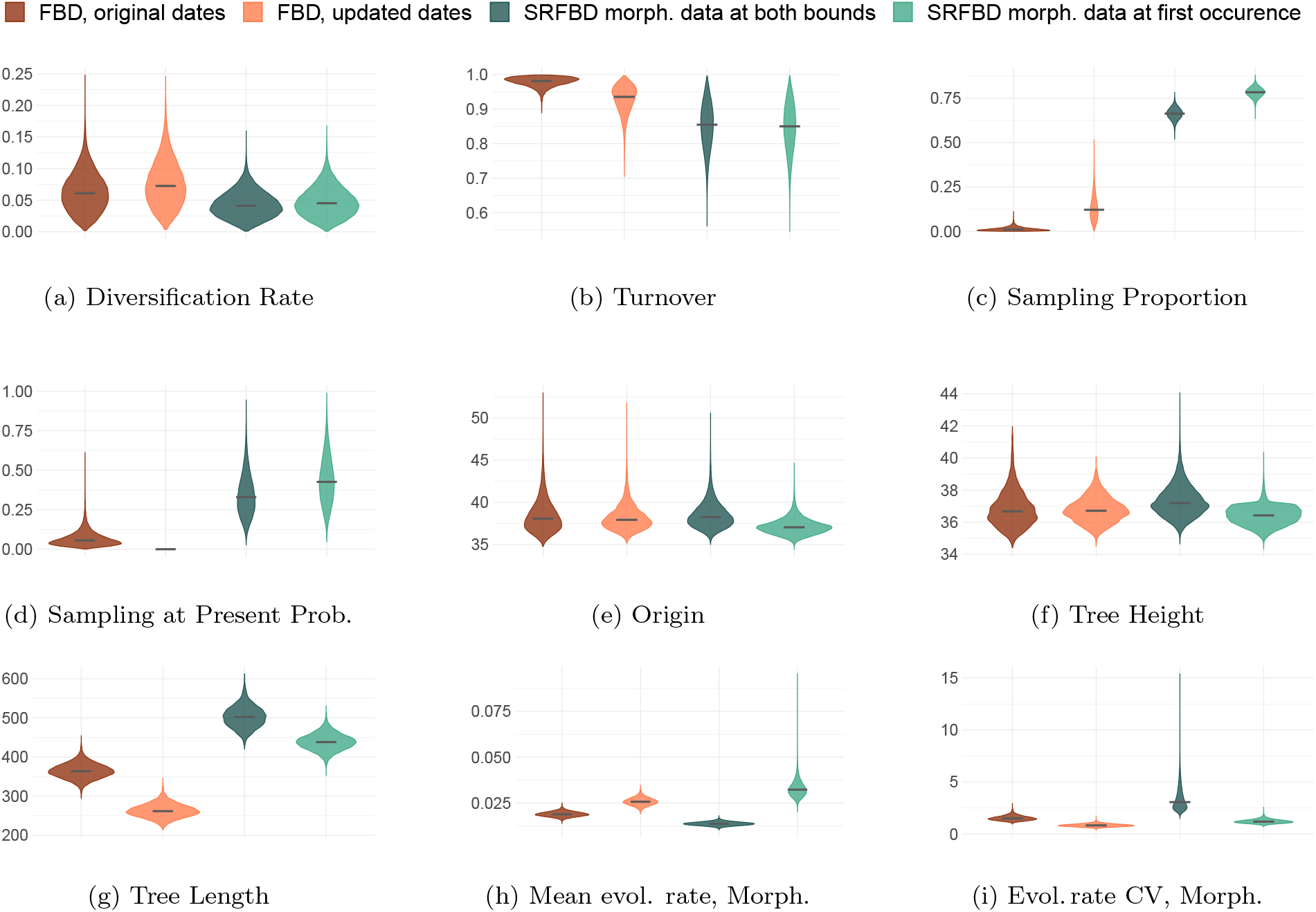
Canid data. Parameter estimates obtained by FBD, FBD with updated dates and SRFBD models, when attaching morphological data to either first occurrence (start of the species range) or both first and last occurrence (both bounds of the species range). SRFBD model always used updated dates. Sampling at present probability (d) was fixed to 0 for FBD analyses with updated dates. The black line shows the median. CV = coefficient of variation.

The morphological evolutionary rate is higher in the SRFBD analysis with morphological data at the first occurrence compared to the FBD analysis, and lower in SRFBD analysis with morphological data at both ends. The coefficient of variation in evolutionary rates for the morphological data is slightly higher in the SRFBD_first_ analysis compared to the FBD analysis, and more than three times higher in the SRFBD_both_ compared to the FBD analysis (Figure 8 (i) and Supplementary Table A.7).

Similarly to the penguin analyses, clade age estimates are generally older in the SRFBD analyses compared to the FBD,(Figure 7 (c) and (d)). When comparing clades that were identified within both FBD and SRFBD_first_ analyses and have a support of 0.2 or higher in either analysis (206 clades in total, Supplementary Figure A.17), the SRFBD_first_ model estimates clade ages to be on average 1.31 Ma older than the FBD model. SRFBD_both_ model estimated clade ages to be on average 0.95 older than FBD (based on 172 clades with the same inclusion criterion, Supplementary Figure A.18). There is also difference in the clade supports. Once morphological data is placed on both ends of stratigraphic ranges in SRFBD analysis, clade supports become closer to the FBD analysis compared to the SRFBD analysis with only first occurrences having morphological data: the mean difference in clade support probability between the models is 0.16 for FBD vs SRFBD_both_, and 0.29 for FBD vs SRFBD_first_ (again when comparing clades with supports of 0.2 or higher in either model).

All four analyses produce very similar estimates for the age of the MRCA of North American canids, with median estimates ranging from 36.42 Ma (SRFBD_first_) to 37.16 Ma (SRFBD_both_). For comparison, in the original analysis [48] with an outgroup and a non-mechanistic tree prior model, the median estimate was 40.76 Ma (95% HPD: [40.4, 41.834]).

Given the small number of morphological characters compared to the number of species, and large amount of missing data, the posterior distributions of trees have large uncertainties. The estimated topological relationships in the SRFBD analysis with morphological data at the first occurrences only is more uncertain than when the data is placed on both ends: the posterior distribution of trees in the SRFBD_first_ analysis contains 3243 clades (in 50K sampled trees) with 342 having posterior supports greater than 0.05 compared to 229 out of 1182 (in 100K sampled trees) in the SRFBD_both_ analysis. With 121 taxa, the most certain distribution (having a single tree) would have 120 clades in the 95% credibility set.

Interestingly, in the tree that has the highest posterior probability, most of the divergences occur within stratigraphic ranges (Figure 9). The 95% HPD intervals for the number of SA species overlap for SRFBD and FBD analyses (Table 2). However, the median number of SAs in the SRFBD_first_ analysis is much higher, at 47, compared to 33 in the FBD. Further, SRFBD_first_ identifies 29 species with probability of being a SA higher than 0.8, while the FBD finds none (Figure 5(b)). Finally, we obtain even more informative (narrower) posterior distributions for the fossil ages than in the penguin analyses (Supplementary Figure A.13a (b)).

**Figure 9:**
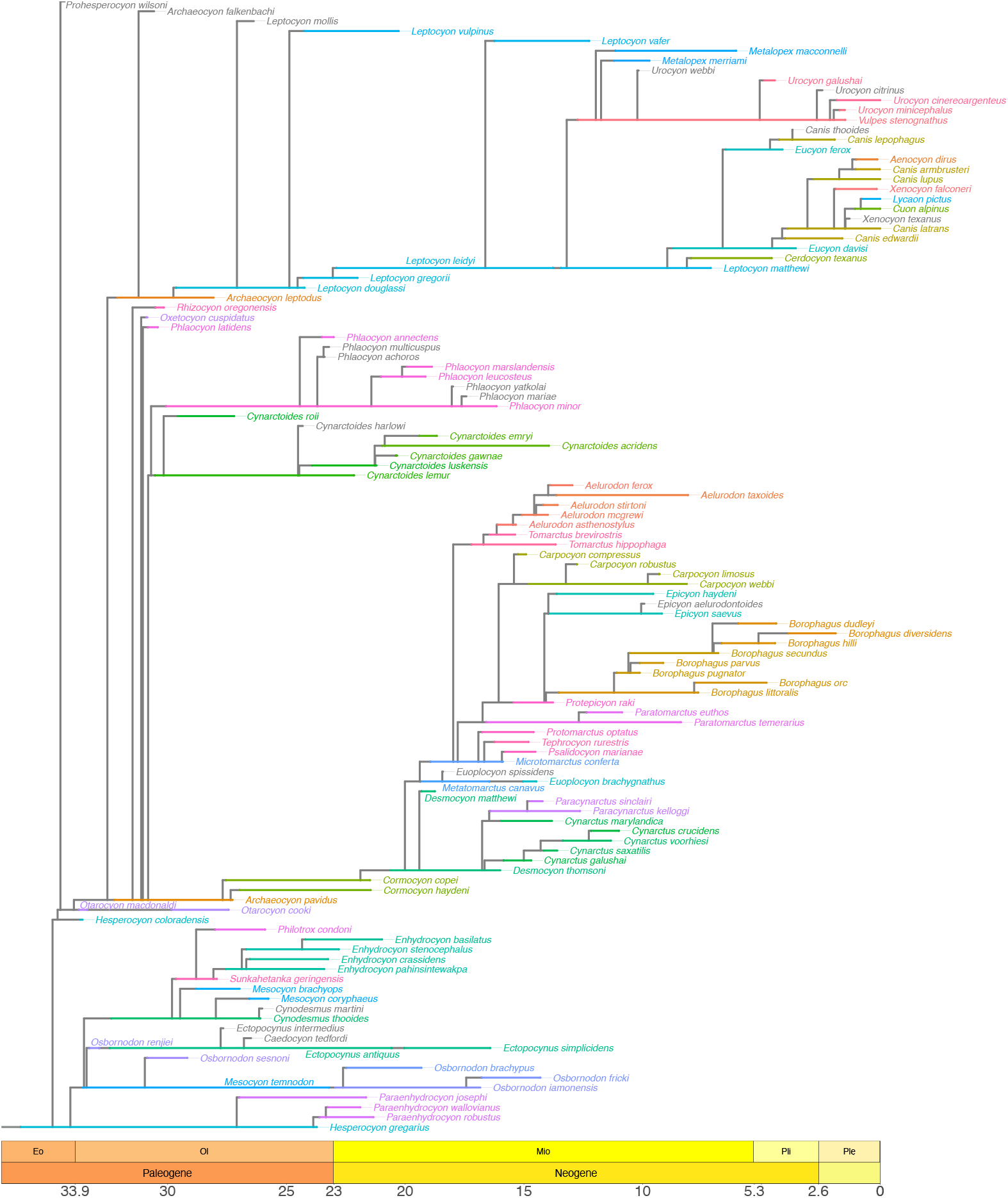
Tree with the maximum posterior probability value for canid data. Non-single occurrence ranges are colored. Time in millions of years before the present (Ma).

## 4 Discussion

Our study shows that accurately representing the stratigraphic ranges of species within phylogenetic inference can produce different, and potentially more accurate, estimates of dated phylogenetic trees and evolutionary parameters, and therefore can influence macroevolutionary interpretations. Our analyses of the penguin and canid families demonstrate that phylogenetic inference using the SRFBD model produces tree topologies, branch lengths and sampled ancestor counts that differ from those obtained using a simpler FBD model. Moreover, the estimated stratigraphic range phylogenetic trees can be used to generate and test new hypotheses, particularly around ancestor-descendant relationships between pairs of sampled species, in comparison with traditional FBD trees. As such, when fossil age information contains stratigraphic ranges, we advocate for the use of the SRFBD model over the simpler FBD for inferring phylogenies.

An important improvement of the SRFBD model over the FBD model is a cleaner estimate of the sampling proportion, due to more accurate modelling of the sampling process. In SRFBD_first_ analyses, we estimated that the analysed fossils represent 42% (median estimate, [30%, 54%] 95% HPD) of extinct penguin species, while the 121 analysed North American canids represent 78% (median estimate, [73%, 84%] 95% HPD) of all extinct canid species. The penguin analysis included 36 fossil species, which implies we estimate there to have been between 67 and 120 extinct species (median 86). There are nearly 70 currently known extinct penguin species [34], which falls within our estimated range. The upper bound estimate of 120 also does not seem unlikely, since new species are still being identified [34, 53]. The fossil record of canids is represented by about 140 [48] known species, with 121 included in this analysis. Our estimated credible interval for the sampling proportion implies that there have been between 144 and 165 (median 155) canid species, meaning that between 4 and 25 are yet to be discovered. The traditional FBD model estimated substantially lower sampling proportions. However, it uses only first occurrence for each species and when we account for the number of known and discarded occurrences (as described in the results section), the FBD estimates come close to those of SRFBD. Using the SRFBD therefore produces a fair estimate of the sampling proportion without requiring an estimate of the number of available fossils.

We do not see large differences in the estimated age of the MRCA in either of the empirical datasets, between the four analyses that we performed. However, for both penguins and canids, using the SRFBD model generally pulled the ages of the internal nodes towards the past: the median age of the penguin crown is estimated to be almost 3 My older (11.25 vs 14.32 Ma) when stratigraphic ranges are modelled, while clades were on average 1.6Ma older for penguins, and 1.31Ma older for canids. As such, using stratigraphic ranges appears to generally push speciation events earlier in the history of a clade. This is likely because the phylogeny needs to accommodate the provided stratigraphic ranges within the branch lengths inferred.

Using the SRFBD has a notable effect on the inference of sampled ancestors. The implemented SRFBD model assumes a budding mode of speciation, and provides a direct and intuitive way of determining which species are ancestral to others. It is possible to infer the directionality of an ancestor-descendant relationship whenever a branching event occurs within a range, or when the ancestral species is a sampled ancestor with a stratigraphic range and the divergence occurs on the continuing lineage. Further, allowing for stratigraphic ranges permits extant species to be inferred as direct sampled ancestors; in the FBD, a choice must be made as to whether extant species are represented as being sampled in the present (at time zero, and therefore cannot be a sampled ancestor) or when a fossil or ancient DNA is available, sampled in the past (permitting being a sampled ancestor, but leaving the model uncertain as to whether the species remains extant). The impact of these model differences is clear but inconsistent in our empirical examples. For the penguins, we observe a decrease in the number of sampled ancestors when using the SRFBD_first_ model compared to the FBD model (medians of 13 and 19 respectively), but the opposite is true for the canids (medians of 47 and 35 respectively). However, all of these estimates are uncertain, with 95% wide HPD intervals that overlap substantially.

An outstanding methodological question is the most appropriate way to attach morphological characters to stratigraphic ranges. In FBD analyses, the morphological change that leads to recognising a given new species has to accumulate prior to the single sampling event of that species, but can be spread along the branch prior to this event. In SRFBD analyses, all morphological changes have to happen before the first occurrence, which is almost always earlier than in the FBD case. When the morphological data are attached to the first occurrence, further morphological evolution can happen later along the branch in this case. However, when the morphological data is placed on both ends, no other changes (apart from possibly a few which must subsequently be reversed) happen for the duration of the range. In our empirical examples, placing morphological data at both ends of the ranges decreased the mean morphological evolutionary rate and increased the variation in morphological evolutionary rates among branches. This can be explained by the fact that the morphological characters are forced to remain unchanged (apart from changes which are later reversed) along the stratigraphic ranges, resulting in the overall smaller evolutionary rate. At the same time, a larger portion of the evolutionary change in morphology is shifted to branches which do not have a stratigraphic range, resulting in the larger variance. Interestingly, we also see that the posterior distribution of trees in the SRFBD analyses is closer (based on clade comparison analysis) to the distribution obtained using the FBD model when morphological data is placed on both ends of the range, compared to only on the first occurrence.

This work enables the community to make full use of the fossil record in combination with molecular data within the Bayesian phylogenetic inference framework BEAST2. While it conceptually enables coherent analysis, we identify several gaps to be addressed in future studies. First, the aforementioned difficulties concerning morphological character attachment to stratigraphic ranges highlight the need for more realistic (nuanced) morphological evolution models. Newly implemented substitution models could average character evolution across the duration of stratigraphic ranges, or model morphological evolution in a manner more similar to punctuated equilibrium, ensuring that character changes take place outside of periods over which conceptually fixed species exist [54, 55]. Alternatively, instead of extending the morphological evolution model, a midpoint sample could be placed within the range purely for the purpose of attaching morphological data to it; this would ‘only’ require changes to the MCMC operators. A second direction for future work is developing methods to better summarise distributions of stratigraphic range phylogenetic trees. In this study, we explored the posterior distributions of the stratigraphic range trees by examining posterior clade heights and supports. Still, in order to visualize a tree, we had to use the maximum posterior probability tree. Developing methods that would produce an analogue of a maximum clade credibility or a centroid tree [56, 57] or use conditional clade distributions to find a tree with highest probability [58] would allow for a more reliable point estimate of a stratigraphic range phylogenetic tree. Third, the SRFBD method will be applicable more broadly once speciation and extinction rates are permitted to vary through time, such as through the inference of piecewise-constant population dynamics [59, 60]. This would not only enable more versatile macroevolutionary analyses, but also allow us to apply the model to epidemiological data, where varying sampling schemes and interventions require time-dependent parameters. Similarly, type-dependent speciation, extinction and sampling rates [61, 62] could be introduced to account for rate heterogeneities due to geography or species traits. Fourth, while our method integrates over speciation events along lineages, it is possible to extend the inference algorithm itself to explicitly estimate them. In particular, a stochastic mapping approach [63, 64] could be used to efficiently impute such events. This would directly allow for the inference of posterior probabilities for direct or indirect ancestry between sampled taxa. Finally, extending the inference algorithm to accommodate the mixed speciation mode version of the model [10] – allowing for asymmetric, symmetric and anagenetic speciation – would allow us to test a whole variety of speciation-related hypotheses.

## Conclusions

Here we present a BEAST2 implementation of the SRFBD model, which permits integration with the available suite of molecular and morphological evolutionary models. The model allows phylogenies to be inferred using a budding mode of speciation, with fossil age information represented as stratigraphic ranges: these novelties make better use of available fossil data and allow for the inference of more biologically realistic phylogenies. We present two empirical case studies, which demonstrate the potential of the model to change how we infer sampled ancestors, as well as produce different estimates of tree topologies and branch lengths compared to the traditional FBD model. In future, further model realism can be added, such as other speciation modes, punctuated evolution models, and evolutionary rate variation across space and time. We envision this framework will enable the testing of hypotheses around ancestor-descendant relationships, and the quantification of macroevolutionary dynamics.

## Acknowledgments

U.S., T.G.V., T.S. and B.J.A. thank ETH Zürich for funding. U.S., T.G.V. and T.S. received funding from the European Research Council (ERC) under the Horizon 2020 Research and Innovation Programme Grant Agreement of the European Union No. 101001077 (PhyCogy). A.G. was supported by Ministry of Business, Innovation, and Employment of New Zealand through an Endeavour Smart Ideas grant (UOOX1912). B.J.A. was supported by Deutsche Forschungsgemeinschaft Collaborative Research Centre 1211 (grant no. 268236062).

The authors thank Joëlle Barido-Sottani for help with the SRFBD tree plotting code.

## Author Contributions

A.G. developed the MCMC kernel. A.G. and U.S. developed the sRanges package and performed data analyses. U.S. performed simulation studies.

B.J.A. compiled stratigraphic range ages and interpreted the results.

A.G., U.S., T.G.V., T.S., B.J.A. conceptualised the study.

U.S., A.G., B.J.A. wrote the manuscript with support from T.G.V. and T.S.

B.J.A., T.G.V. and T.S. supervised the project.

## Data and Software Availability

See Supplementary Materials of [37] for detail description of morphological data and GenBank accession numbers for molecular data (Table S1 in Ksepka et al. [37]).

See https://github.com/jugne/sRanges-material for BEAST 2.7 XML files, used to run the analyses and scripts used to analyse the data and produce figures.

See https://github.com/jugne/stratigraphic-ranges for the sRanges BEAST2.7 package source code and installation.

## Appendices

### A Supplementary Materials

#### Operators

The operators for orientated trees with stratigraphic ranges need to simultaneously manage orientation at each internal node and stratigraphic information, ensuring that each proposed tree modification respects both budding speciation and geological context. To this end, we introduced several modifications to the classic tree operators.

##### Adjusted Wilson-Balding Operator

This is a version of the Wilson-Balding operator for oriented trees that also abides by stratigraphic range constraints.

1. Node selection: select a non-root node *i* for reposition. This node is not a sampled ancestor (SA) or part of a stratigraphic range.
2. New location identification: identify a new location on a tree for node *i* by selecting another node *j*, that is not a sibling node of *i* or its parent. If *j* is a leaf, it will become SA parent of *i*, then it must be older than *i*. Otherwise, it will become a sibling node to *i* and *j* parent node must be older than *i*.
3. Move execution: detach the subtree, rooted at node *i* and re-attach it. If *j* is a leaf, its height is the new height of *i* parent node. Otherwise randomly choose the height of *i* parent node between height of *j* and its parent node, so that *i* and *j* are now sibling nodes. Attachment place respects existing node orientations and place *i* in a logical temporal position (i.e., not making an ancestor younger than its descendant.
4. Direct proposal rejection: if any step of the move violates the constraints of the oriented tree or stratigraphic logic (e.g., no valid position found for reattachment, invalid temporal positioning), the proposal is immediately rejected, avoiding the new posterior calculation.

##### Adjusted LeafToSampledAncestor Operator

This is a version of operator, previously implemented in Gavryushkina et al. [6], adjusted to respect stratigraphic range constraints. Since creating SA ranges is already ensured by WB operator above, this operator deals only with *single-node ranges*. That is, when species is represented by a single node, not a range with temporally distinct bounds.

1. Node selection: The operator selects a node *i* (either a leaf or a SA node) that is a part of single-node range.
2. Move execution:
  a. if *i* is a leaf node, the operator turns it into a SA node by first selecting a non-range edge with start older than node *i* and end more recent than node *i*. Then, it splits this edge, by placing node *i* on it. Node *i* becomes a degree two node).
  b. if *i* is a SA node, it is transformed to leaf node by choosing a new parent node height. Node *i* becomes a degree one node.
3. Orientation: the operator accounts for random orientation assignment when going from SA node to a leaf node.
4. Direct proposal rejection: if any step of the move violates the constraints of the oriented tree or stratigraphic logic (e.g., no valid position found for reattachment, invalid temporal positioning), the proposal is immediately rejected, avoiding the new posterior calculation.

##### LeftRightChildSwap Operator

This operator swaps the orientation of children nodes for an internal tree node that is not a SA and not a node lying on a straight line between a species range bounds.

## Supplementary figures

**Figure A.10:**
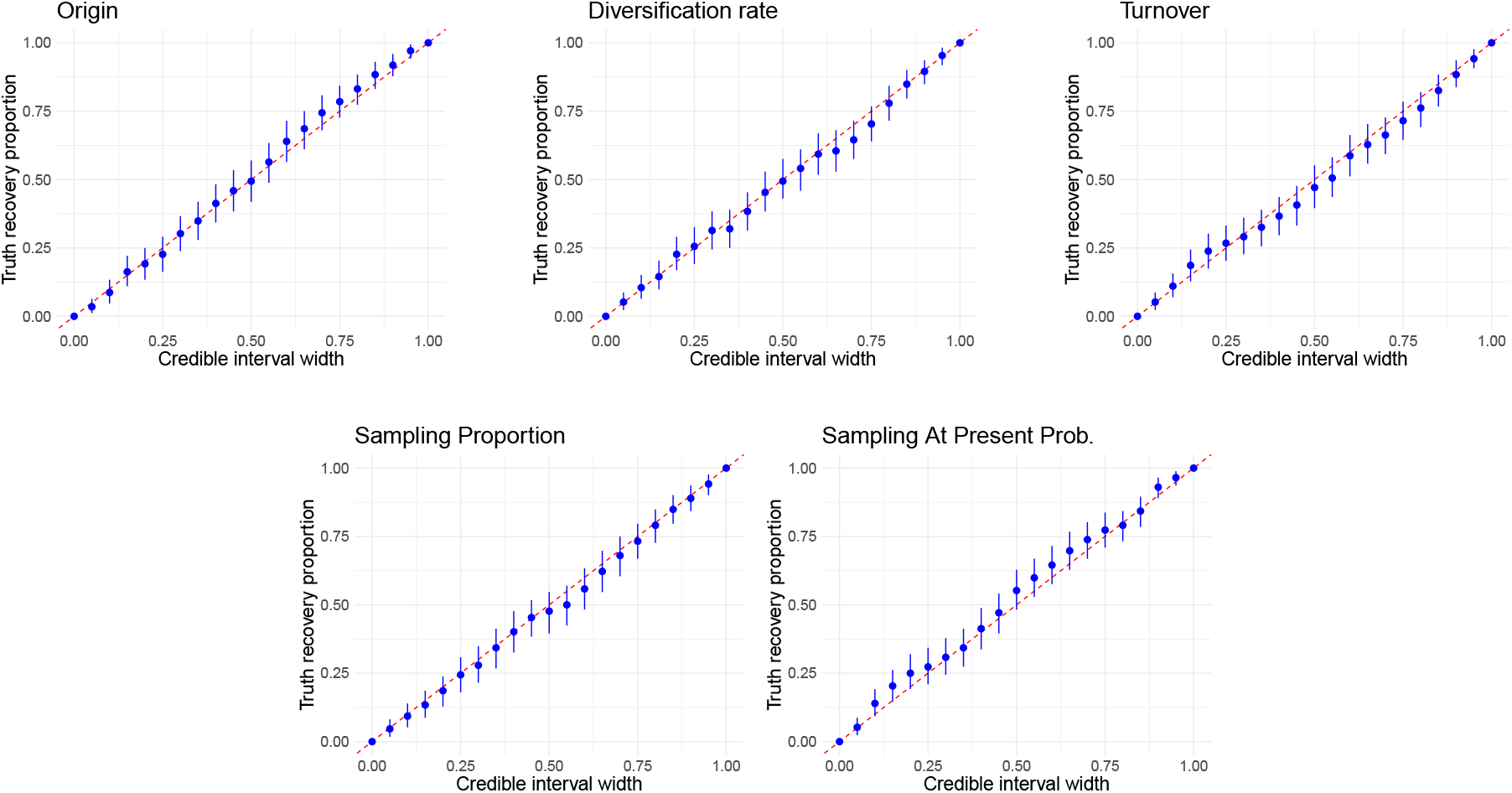
Results of the validation simulation study, molecular data attached to all included samples. Blue dots show the fraction of analyses where the true value is within the HPD credible interval of a certain width. The blue vertical lines show bootstrap confidence intervals. The diagonal red dashed line *x* = *y* is the expected ideal result.

**Figure A.11:**
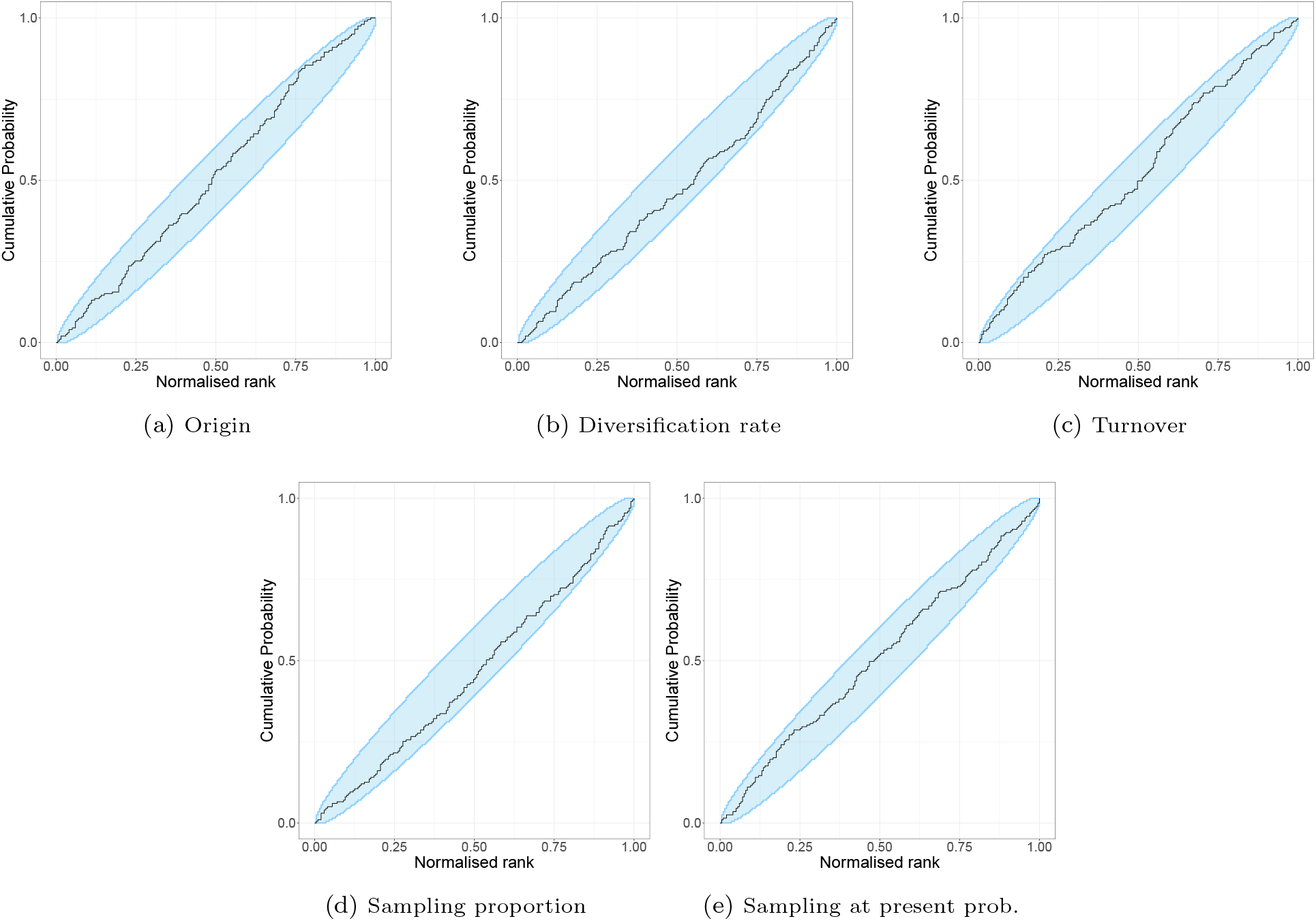
Results of the rank-uniformity validation, molecular data attached to extant samples. The empirical cumulative distribution function (ECDF). Light-blue areas show confidence inference about the ECDF. Whenever the ECDF falls within this confidence interval, we say that the rank is uniform and validation is passed [17, 18].

**Figure A.12:**
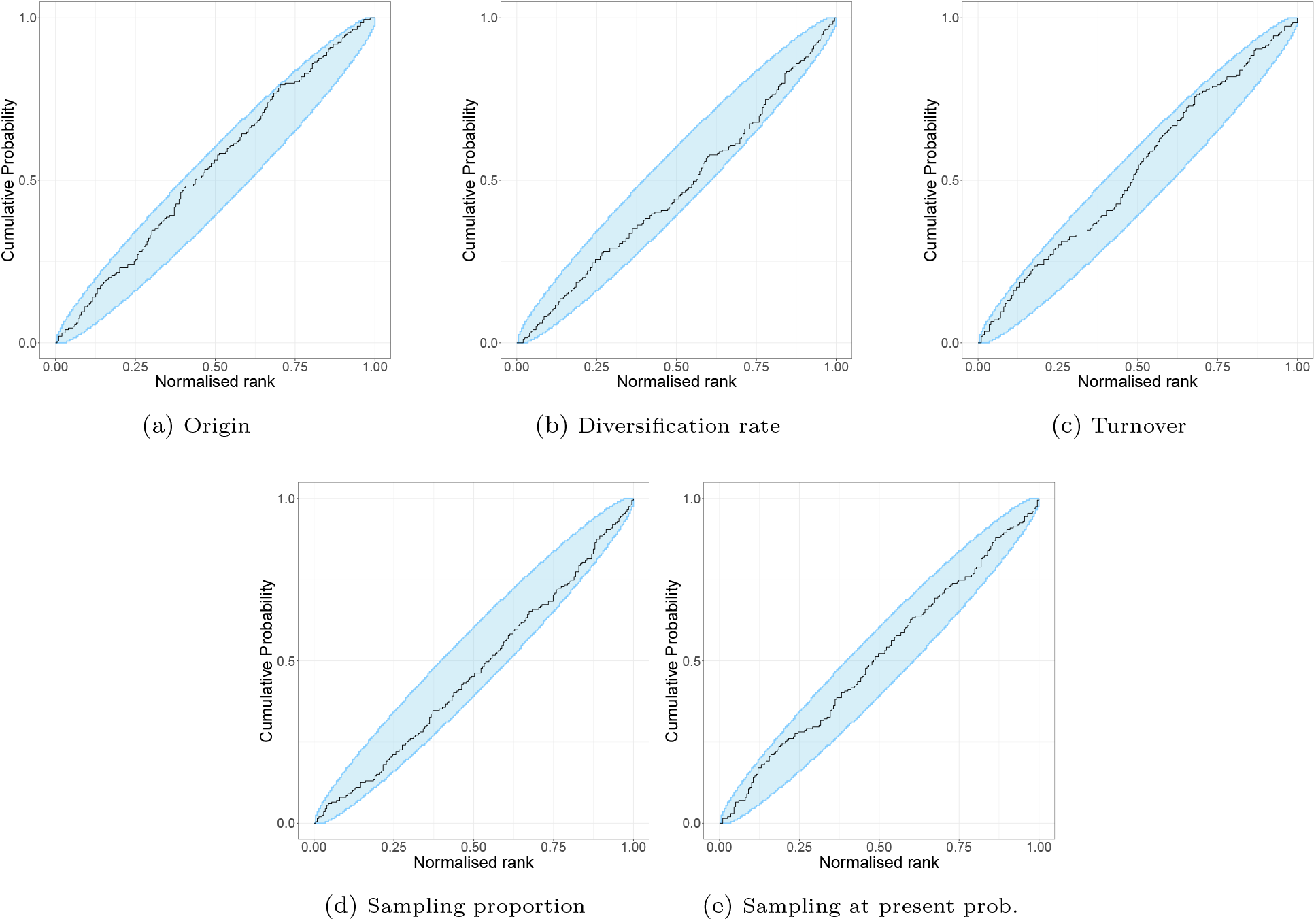
Results of the rank-uniformity validation, molecular data attached to all included samples. The empirical cumulative distribution function (ECDF). Light-blue areas show confidence inference about the ECDF. Whenever the ECDF falls within this confidence interval, we say that the rank is uniform and validation is passed [17, 18].

**Figure A.13:**
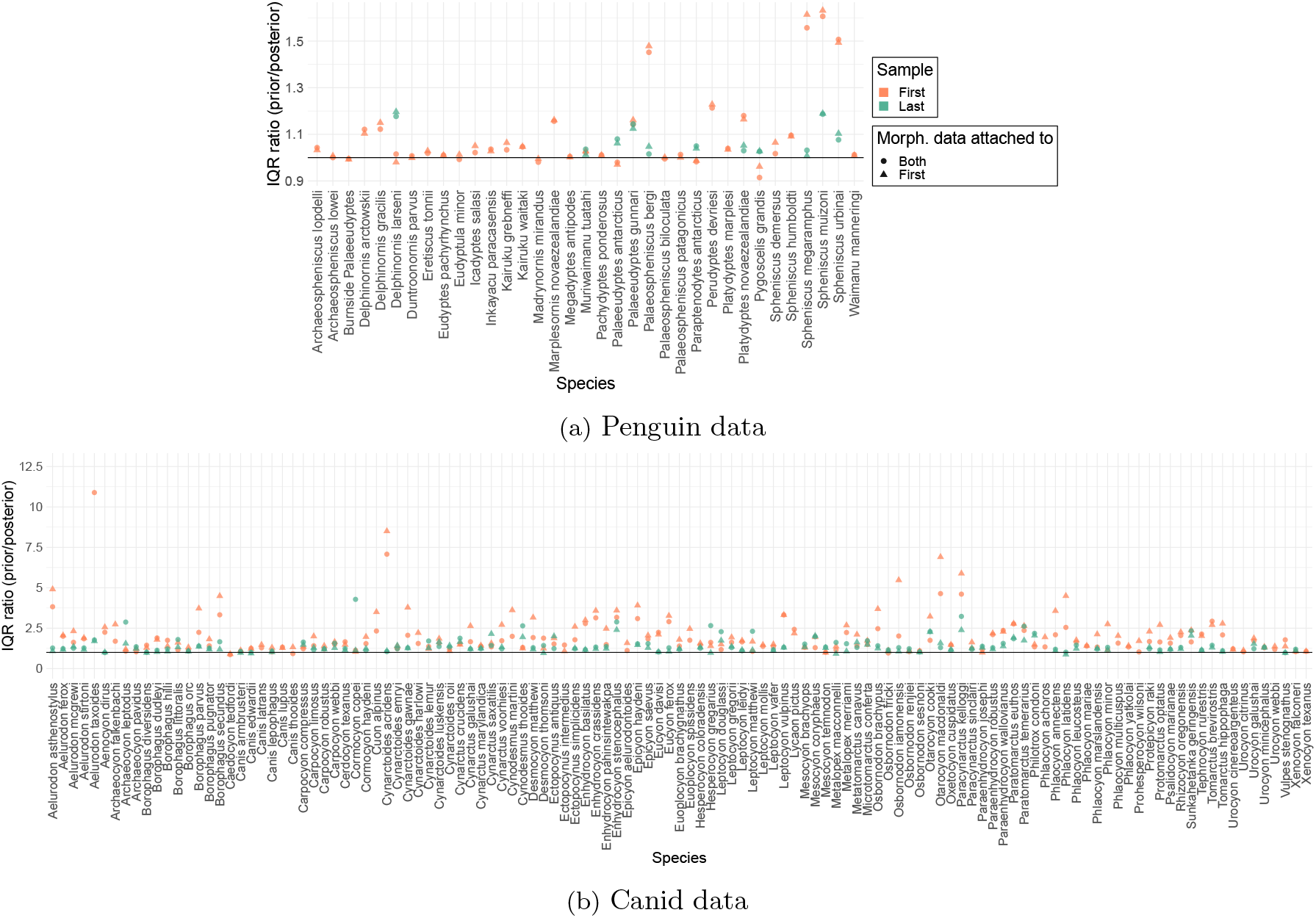
Ratio of the interquantile range (IQR) between the uniform prior and posterior of first and last species sample by SRFBD. Dots show analyses when morphological data was attached to both samples and triangles when it was attached only to the first sample. The black horizontal line shows a ratio of 1 (IQR of prior and posterior are equal). The higher value is above the line, the more narrow and informative is the posterior compared to the prior.

**Figure A.14:**
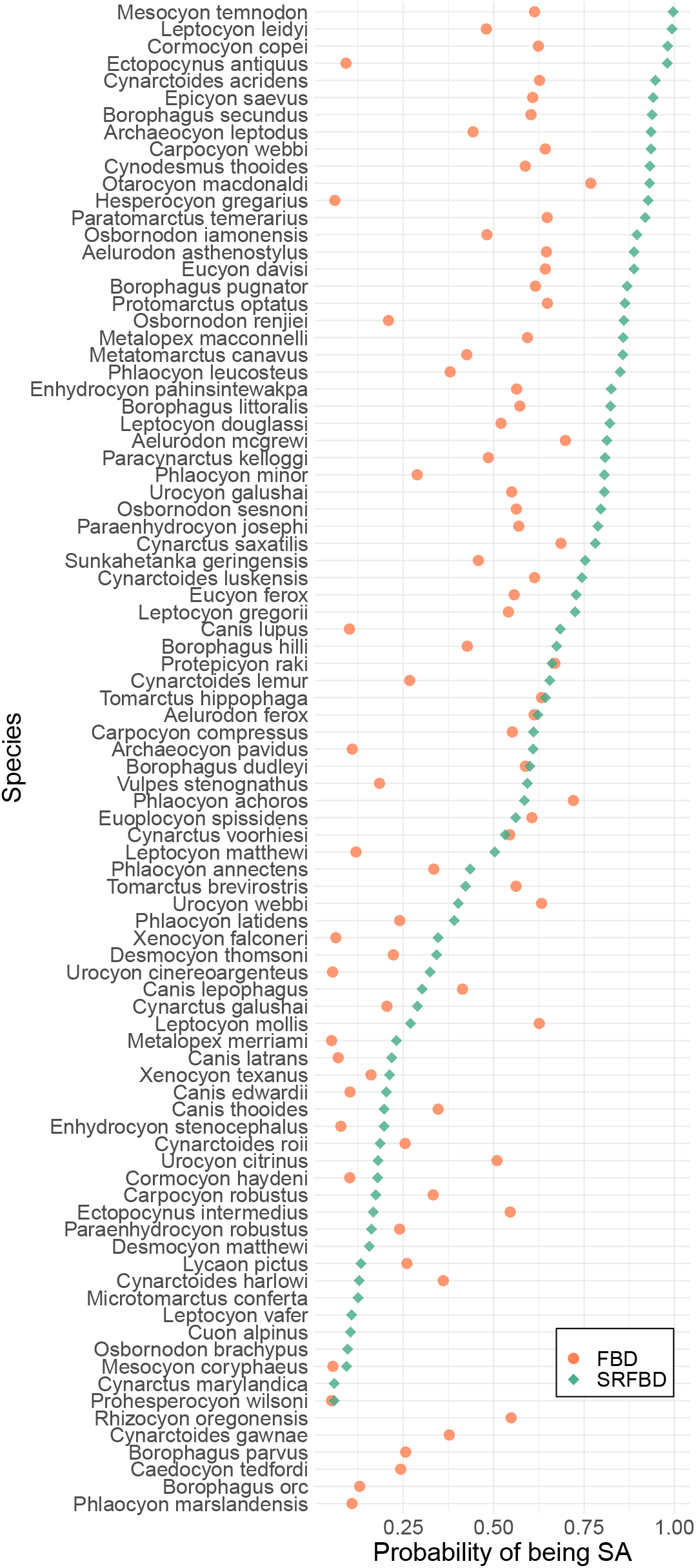
Canid data. Posterior probability of being a SA species. Circles for results by FBD, and diamonds for results by SRFBD_first_.

**Figure A.15:**
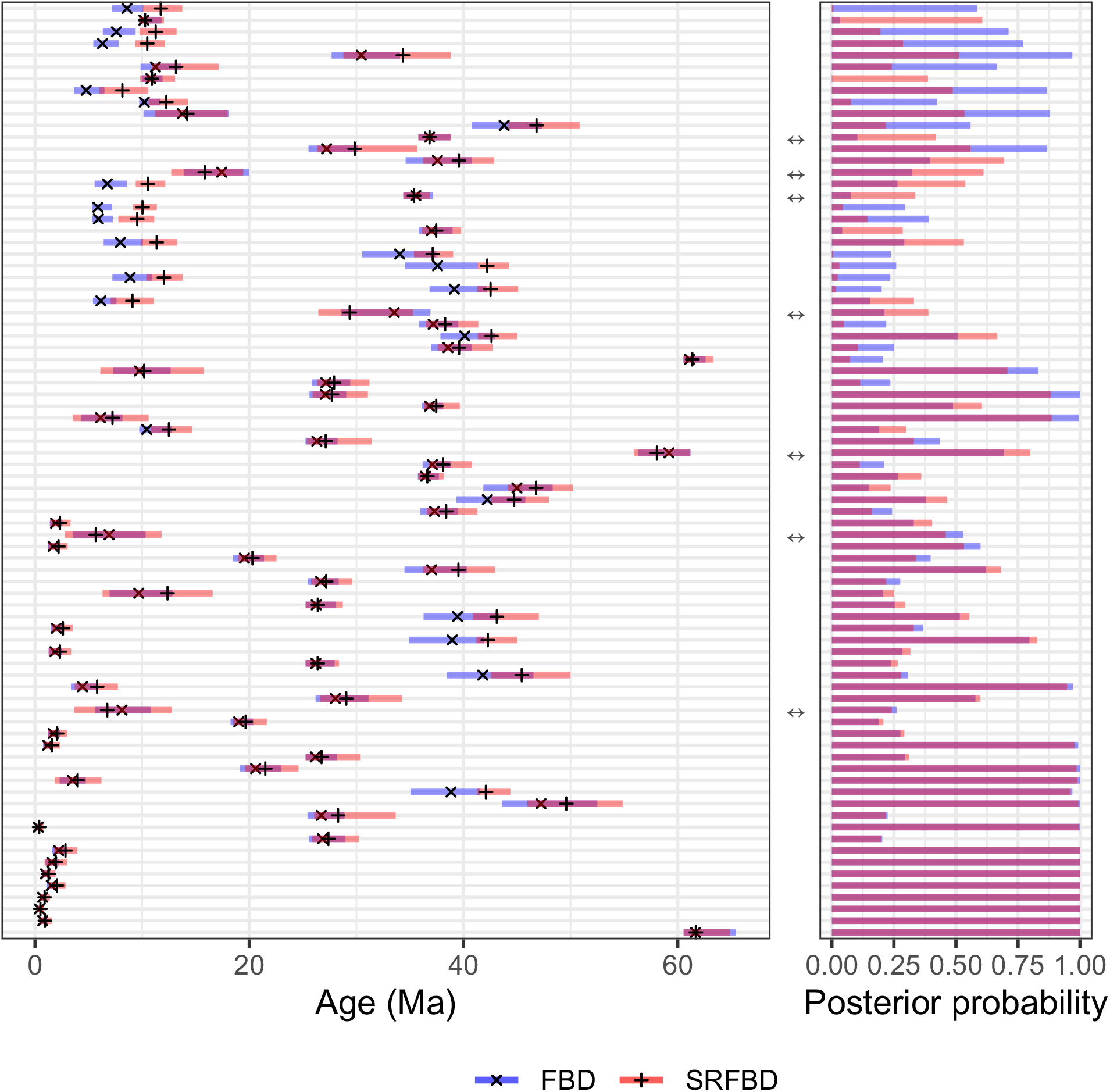
Penguin data. **Left** Mean ages and 95% HPD intervals identified with SRFBD_first_ and FBD. Means are marked by *×* (FBD) and + (SRFBD). **Right** Posterior clade support. Both plots are aligned so that the same clade is represented on y axis. Only clades that were identified in both analyses and had posterior support of 0.2 or higher in at least one of them are included. Double arrows between the plots note the clades for which mean age is estimated older by FBD, compared to SRFBD. Total included clades: 80. Clades, for which mean age is estimated older by FBD: 7. Clades for which the 95% HPD interval was estimated more narrow by SRFBD than FBD: 17.

**Figure A.16:**
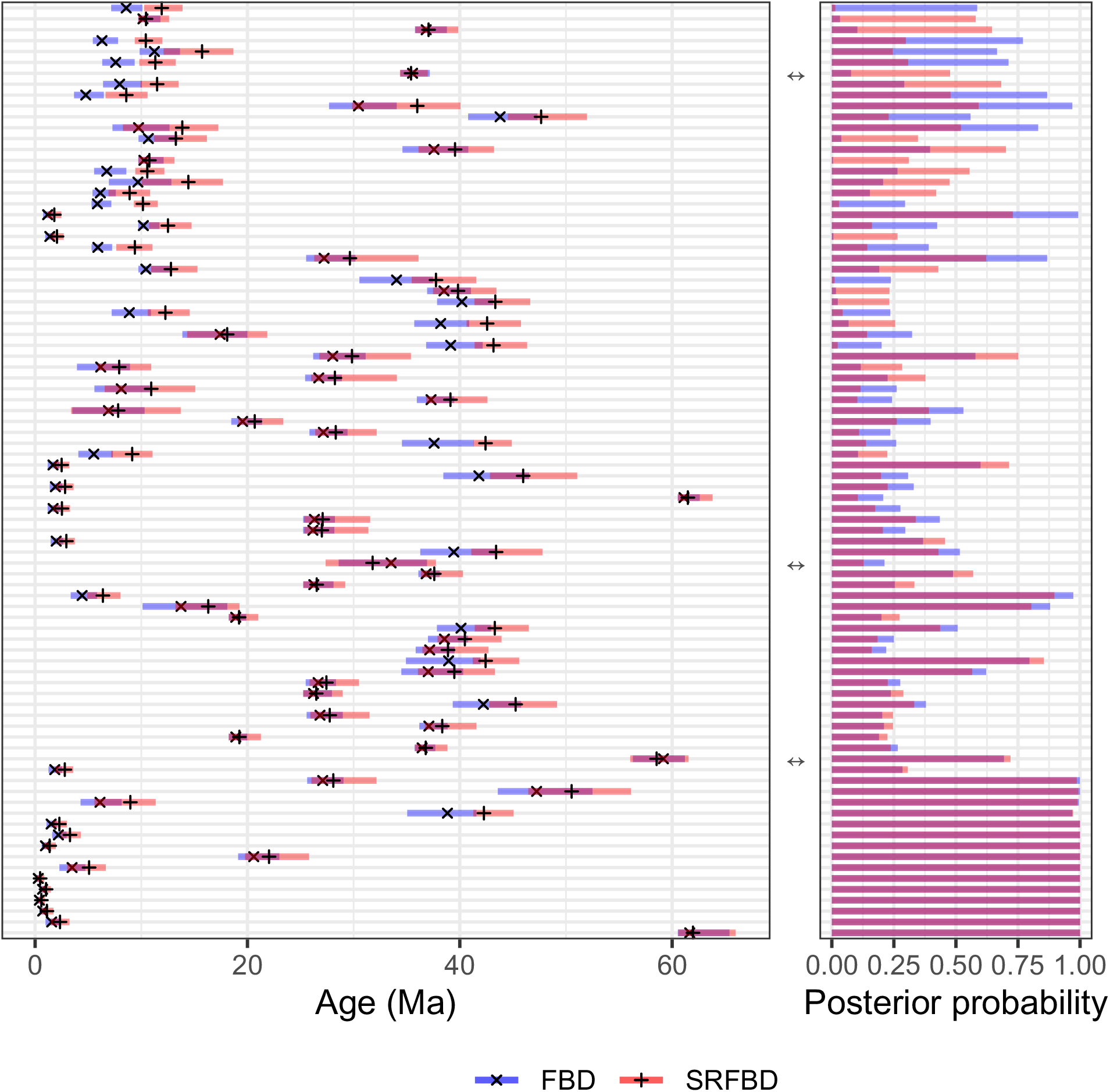
Penguin data. **Left** Mean ages and 95% HPD intervals identified with SRFBD_both_ and FBD. Means are marked by *×* (FBD) and + (SRFBD). **Right** Posterior clade support. Both plots are aligned so that the same clade is represented on y axis. Only clades that were identified in both analyses and had posterior support of 0.2 or higher in at least one of them are included. Double arrows between the plots note the clades for which mean age is estimated older by FBD, compared to SRFBD. Total included clades: 86. Clades, for which mean age is estimated older by FBD: 3. Clades for which the 95% HPD interval was estimated more narrow by SRFBD than FBD: 14.

**Figure A.17:**
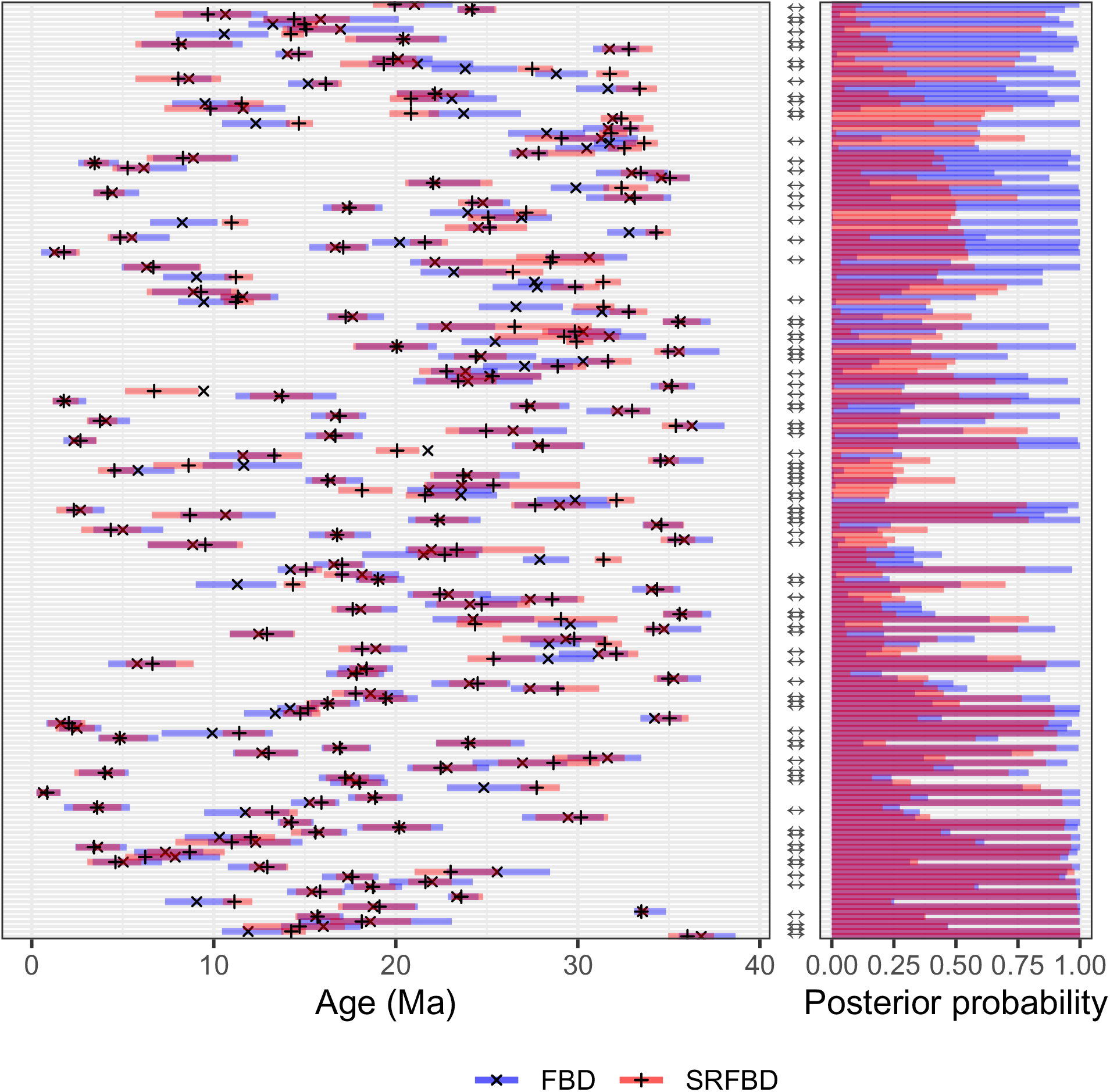
Canid data. **Left** Mean ages and 95% HPD intervals identified with SRFBD_first_ and FBD. Means are marked by *×* (FBD) and + (SRFBD). **Right** Posterior clade support. Both plots are aligned so that the same clade is represented on y axis. Only clades that were identified in both analyses and had posterior support of 0.2 or higher in at least one of them are included. Double arrows between the plots note the clades for which mean age is estimated older by FBD, compared to SRFBD. Total included clades: 189. Clades, for which mean age is estimated older by FBD: 85. Clades for which the 95% HPD interval was estimated more narrow by SRFBD than FBD: 154.

**Figure A.18:**
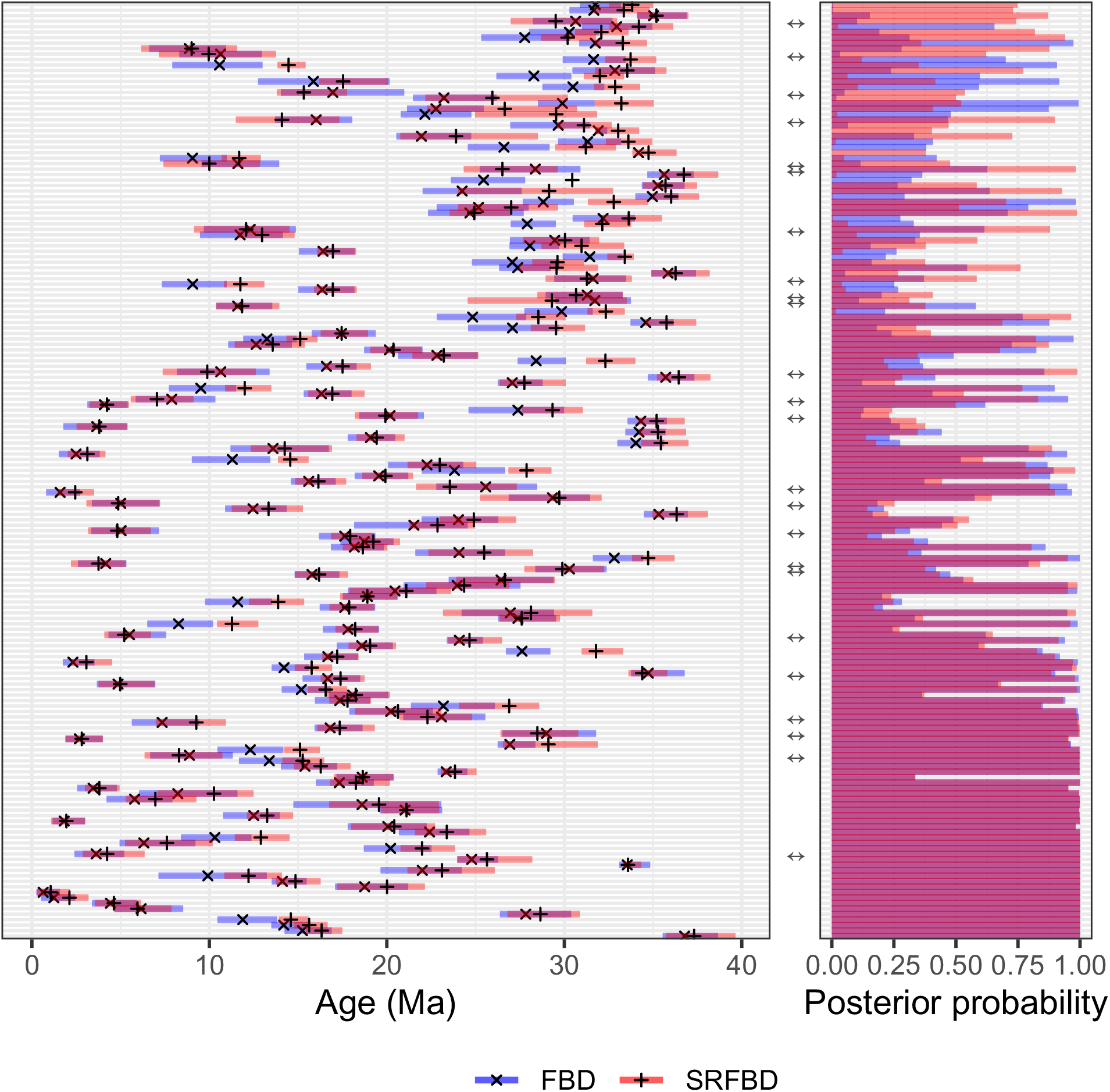
Canid data. **Left** Mean ages and 95% HPD intervals identified with SRFBD_both_ and FBD. Means are marked by *×* (FBD) and + (SRFBD). **Right** Posterior clade support. Both plots are aligned so that the same clade is represented on y axis. Only clades that were identified in both analyses and had posterior support of 0.2 or higher in at least one of them are included. Double arrows between the plots note the clades for which mean age is estimated older by FBD, compared to SRFBD. Total included clades: 171. Clades, for which mean age is estimated older by FBD: 24. Clades for which the 95% HPD interval was estimated more narrow by SRFBD than FBD: 88.

## Supplementary tables

**Table A.5:**
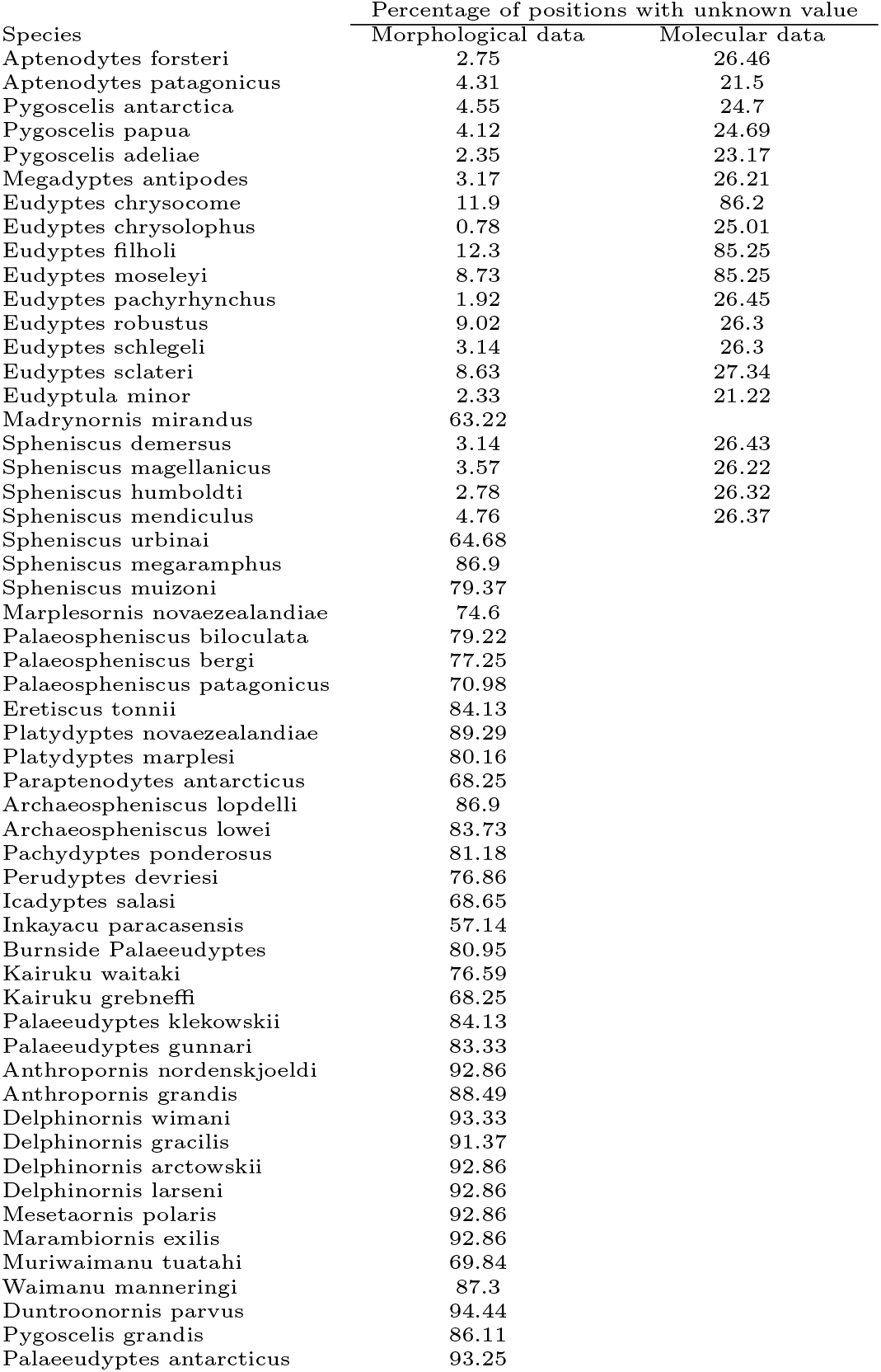
Penguin data. Percentage of missing data for morphological and molecular data (when available).

**Table A.6:**
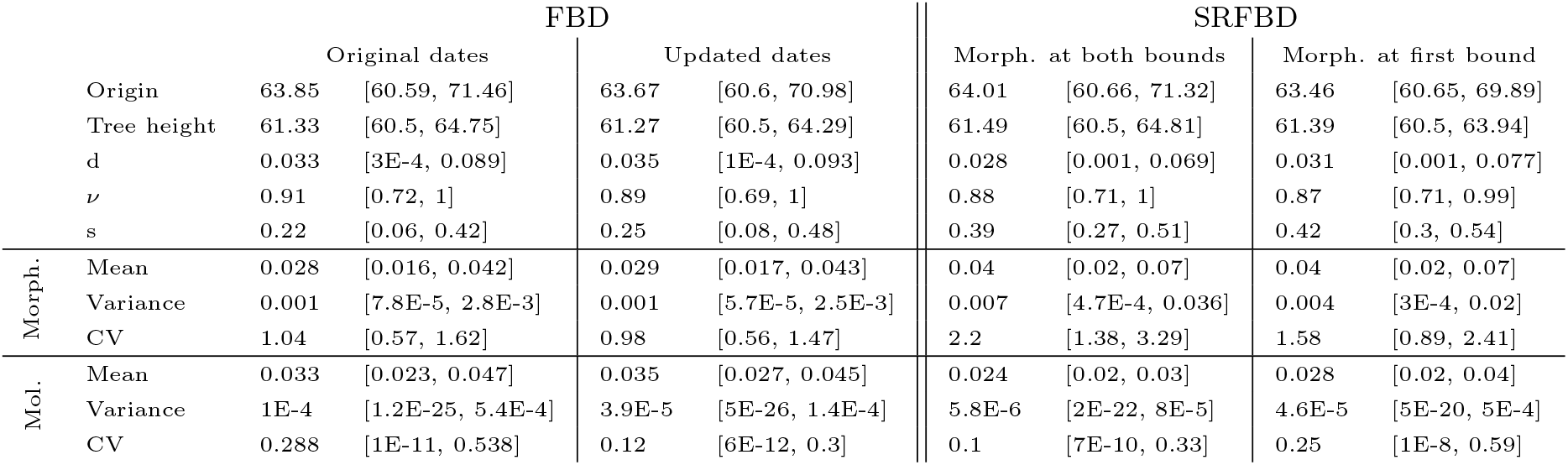
Penguin data. Median tree statistics and parameter estimates obtained by FBD and SRFBD. CV=coefficient of variation. Morphological (Morph.) and molecular (Mol.) evolutionary rates measured in substitutions per site per million years. 95% HPD intervals in the square brackets.

**Table A.7:** Canid data. Median parameter estimates obtained by FBD and SRFBD. CV=coefficient of variation. Morphological (Morph.) and molecular (Mol.) evolutionary rates measured in substitutions per site per million years. 95% HPD intervals in the square brackets.

|  |  | FBD |  |  |  | SRFBD |  |  |  |
| --- | --- | --- | --- | --- | --- | --- | --- | --- | --- |
|  |  | Original fixed dates |  | Updated dates |  | Morph. at both bounds |  | Morph. at first bound |  |
|  | Origin | 38.06 | [35.24, 42.45] | 37.92 | [35.68, 41.25] | 38.19 | [35.87, 41.56] | 37.03 | [35.4, 39.06] |
|  | Tree height | 36.68 | [34.93, 39.05] | 36.71 | [35.24, 38.28] | 37.16 | [35.46, 39.19] | 36.42 | [35.14, 37.32] |
|  | d | 0.061 | [7.13E-3, 0.12] | 0.073 | [0.012, 0.146] | 0.041 | [5.66E-3, 0.083] | 0.045 | [5.87E-3, 0.092] |
| | $\nu$ | 0.98 | [0.95, 1] | 0.94 | [0.85, 0.99] | 0.86 | [0.73, 0.97] | 0.85 | [0.71, 0.97] |
|  | s | 0.01 | [2E-4, 0.03] | 0.12 | [0.01, 0.26] | 0.66 | [0.6, 0.72] | 0.78 | [0.73, 0.84] |
| | $\rho$ | 0.06 | [4.4E-3, 0.16] | 0 | - | 0.33 | [0.11, 0.6] | 0.43 | [0.16, 0.73] |
| Morph. | Mean | 0.019 | [0.016, 0.022] | 0.026 | [0.022, 0.03] | 0.014 | [0.012, 0.016] | 0.032 | [0.025, 0.042] |
|  | Variance | 2.3E-3 | [0.75E-3, 5.56E-3] | 0.66E-3 | [0.2E-3, 1.41E-3] | 0.012 | [1.3E-3, 0.09] | 2.3E-3 | [0.7E-3, 5.5E-3] |
|  | CV | 1.5 | [1.11, 1.97] | 0.83 | [0.59, 1.11] | 3.08 | [1.9, 5.72] | 1.19 | [0.88, 1.59] |

## Footnotes

1 As there is no conventional terminology for distributions in hierarchical models, we call this tree distribution a prior, while noting it could also be considered a likelihood for *η*.

## Notes

https://github.com/jugne/sRanges-material

https://github.com/jugne/stratigraphic-ranges

### Competing Interest Statement

The authors have declared no competing interest.

